# Dietary fiber controls blood pressure and cardiovascular risk by lowering large intestinal pH and activating the proton-sensing receptor GPR65

**DOI:** 10.1101/2022.11.17.516695

**Authors:** Liang Xie, Rikeish R. Muralitharan, Evany Dinakis, Simona Antonacci, Kwan Charmaine Leung, Zoe McArdle, Katrina Mirabito Colafella, Michael Nakai, Madeleine Paterson, Alex Peh, Hamdi Jama, Ekaterina Salimova, Dovile Anderson, Caroline Ang, Md Jahangir Alam, Yu-Anne Yap, Darren Creek, Remy Robert, Joanne A. O’Donnell, Charles R. Mackay, Francine Z. Marques

**Affiliations:** Hypertension Research Laboratory, Victorian Heart Institute and Department of Pharmacology, Biomedical Discovery Institute, Faculty of Medicine, Nursing and Health Sciences, Monash University, Clayton VIC 3800, Australia; Department of Microbiology, Biomedicine Discovery Institute, Monash University, Clayton, VIC 3800, Australia; Precision Medicine Translational Research Programme, Department of Obstetrics & Gynaecology, Yong Loo Lin School of Medicine, National University of Singapore, Singapore; Cardiovascular Disease Program, Biomedicine Discovery Institute and Department of Physiology, Faculty of Medicine, Nursing and Health Sciences, Monash University, Clayton VIC 3800, Australia; Bioimaging Platform, La Trobe University, Melbourne Australia, Monash Biomedical Imaging, Monash University, Clayton, VIC 3168, Australia; Monash Proteomics and Metabolomics Facility, Monash Institute of Pharmaceutical Sciences, Monash University, Parkville, VIC 3052, Australia; Department of Physiology, Biomedicine Discovery Institute, Monash University, Clayton, VIC 3800, Australia; School of Pharmaceutical Sciences, Shandong Analysis and Test Center, Qilu University of Technology (Shandong Academy of Sciences), Jinan, 250014, China; Baker Heart and Diabetes Institute, Melbourne, VIC 3000, Australia

**Keywords:** hypertension, metabolites, microbiome, pH, G-protein coupled receptor

## Abstract

High blood pressure (BP) is the most common cause of death globally, due to increasing the risk of cardiovascular diseases. Dietary fiber regulates BP through gut microbial production of acidic metabolites known as short-chain fatty acids (SCFAs). The specific mechanisms of how SCFAs regulate BP are still emerging. In a phenome-wide association study, we identified that the proton-sensing G-protein-coupled receptor *GPR65* gene is associated with hypertension and its associated end-organ damage phenotypes. We hypothesized that acidic metabolites produced from the gut microbiota may activate GPR65, thus conferring BP regulating effects. We found that dietary fiber levels determined the luminal and interstitial tissue pH in the large intestine through production of SCFAs by the gut microbiota. We identified that low pH produced by high fiber intake, acting via GPR65 signaling, increased cAMP production and phosphorylation of CREB, and restricted the production of hypertension-promoting inflammatory cytokines by CD8^+^ T cells. *Gpr65^−/−^* mice spontaneously developed higher BP, cardiac and renal hypertrophy and fibrosis. We showed that the benefits of a diet high in fiber, which prevented hypertension and associated end-organ damage, were decreased in *Gpr65^−/−^* mice. Finally, adoptive transfers revealed that GPR65 deficiency in CD8^+^ T cells causally explained this phenotype. In conclusion, we showed that pH sensing by GPR65 in CD8+ T cells mediates much of the cardiovascular benefits of dietary fiber. pH sensing represents a novel gene-by-environment interaction of gut microbiota-to-host biological effects and may form the basis for new therapeutic strategies for hypertension.

## Introduction

Hypertension accounted for 19.2% of all deaths (10.8 million deaths) worldwide in 2019, primarily due to cardiovascular disease (CVD).^1^ Insufficient intake of foods high in dietary fiber, such as vegetables, fruits and wholegrain foods, contributes to mortality and morbidity of non-communicable diseases^2^ resulting from higher blood pressure (BP) and CVD.^3^ Gut microbial metabolites, called short-chain fatty acids (SCFAs), produced upon dietary fiber fermentation, underlie the cardiorenal benefits conferred by dietary fiber.^4–7^ As a consequence, dietary interventions that increase colonic SCFAs (either via direct administration of SCFAs or a high fiber diet) have been reported to reduce BP significantly not only in experimental hypertensive animals^4,5^ but, more recently, in untreated essential hypertensive patients in a phase II randomized clinical trial.^8^ However, the underlying mechanism of how SCFAs lower BP remains to be fully elucidated. Thus far, studies have focused on the direct role of SCFAs, via binding to and activating G-protein coupled receptors (GPCRs) such as GPR41 and GPR43.^4,5,9,10^

An important observation that has not been explored yet is that the production of SCFAs from the bacterial fermentation of dietary fiber in the colon reduces the luminal pH in the distal small intestine from ∼8 to ∼6.5.^11–13^ Such acidic intestinal pH ∼6.5 created by SCFAs may activate proton-sensing receptors, including the G-protein-coupled receptor GPR65.^14–17^ GPR65 is critical for gut homeostasis. A set of genetic variants (e.g., rs3742704, rs8005161) in the human *GPR65* gene are strongly associated with inflammatory bowel disease (IBD),^18–23^ and GPR65-deficient mice exhibited exacerbated inflammation in experimental colitis models.^23–25^ In addition, the *GPR65* minor T allele of rs8005161 was associated with high BP in a phenome-wide association study (PheWAS).^22^ These genotypes are associated with ∼50% reduced GPR65 signaling,^23^ and provide an insight into the relevance of GPR65 in human disease. We validated these results in an independent PheWAS using the Atlas of GWAS Summary Statistics data, showing that *GPR65* is associated with hypertension, cardiac hypertrophy and function (e.g., ejection fraction, fractional shortening), and heart failure (Table S1).^26^ Moreover, GPR65 is highly expressed by various leukocyte subsets^27–29^ and regulates inflammatory responses relevant to hypertension, such as inhibiting inflammatory cytokine production.^22^ Therefore, proton sensing by GPR65 may be an instrumental link between fiber, colonic luminal pH, and cardiovascular health.

Here we aimed to establish the role of colonic pH and the proton-sensing receptor GPR65 in hypertension and its associated end-organ damage that are risk factors for CVD, particularly when fermentable fiber intake is high. We conducted a series of studies in mice that provided evidence for a novel role of GPR65 in the cardiovascular protection conferred by dietary fiber. We demonstrate that dietary fiber confers cardiovascular protection by lowering colonic pH and activating downstream signaling of the proton-sensor GPR65 that lowers inflammation via inhibiting inflammatory cytokines produced by CD8+ T cells.

## Methods

For detailed description of Methods, please see the Online Supplemental Files.

### Animal studies

All animal care and experimental procedures used in this study were approved by the Animal Ethics Committee of Monash University (17465, 27929, 37720). Wild-type (WT) C57BL/6J mice were obtained from the Monash Animal Research Platform, Monash University, housed in the same facility under the same conditions. *Gpr65^−/−^* mice on a C57BL/6J background were generated by using CRISPR/Cas9-based protocol at the Monash Genome Modification Platform (MGMP) at Monash University. *Gpr65^gfp/gfp^* mice were purchased from The Jackson Laboratory (stock number 008577). *Gpr65^gfp/gfp^* mice have 90% of exon 2 coding sequences replaced by promoterless IRES-EGFP sequences to disrupt GPR65 function.^30^ *Gpr65^gfp/+^* mice were generated by crossing *Gpr65^gfp/gfp^* mice with WT C57BL/6J mice. *Rag1^−/−^* mice on a C57BL/6J background were obtained from the Clive and Vera Ramaciotti Laboratories, Walter and Eliza Hall Institute of Medical Research. All mice were maintained under specific pathogen-free and controlled environmental conditions.

Murine diets used in this study include: i) Normal chow diet (Barastoc rat and mouse pellets; Ridley); ii) AIN93G (Specialty Feeds); iii) a diet lacking resistant starches (low fiber, LF, SF09-028; Specialty Feeds); iv) a diet rich in resistant starches (high fiber with all carbohydrates replaced with resistant starches, HF, SF11-025; Specialty Feeds) (Table S2). To eliminate the gut microbiota, mice were treated with a combination of enrofloxacin (Baytril, 10 mg/kg body weight/ day) and amoxicillin with clavulanic acid (50 mg/kg body weight/ day) in drinking water for a week before the experimental procedures and maintained on this antibiotic cocktail until the endpoint.

### Tissue Collection

Upon euthanasia by CO_2_ asphyxiation, blood and various tissues (heart, kidney, liver, gut, cecal content, spleen) were collected for further analyses. Samples were either snap-frozen using liquid nitrogen and stored at −80 °C for subsequent DNA/RNA extractions, stored in formalin before paraffin embedding for histopathology studies, or kept in cold phosphate buffer saline (PBS) to be used fresh for flow cytometry studies.

### Large intestinal intraluminal and interstitial fluid pH measurement in mouse

The pH was measured in samples collected from the cecum and colon using a pre-calibrated Thermo-Scientific Orion Star A211 pH meter as previously described.^31,32^ Intestinal interstitial fluid pH was measured using a protocol generously shared by Professor Dominik Muller (Max Delbrück Center for Molecular Medicine in the Helmholtz Association, Berlin, Germany), adapted from ^33^. The same Orion PerpHec ROSS Combination pH Micro Electrode (Thermo-scientific, diameter 3mm) was used to measure the pH of the intestinal interstitial fluid. We ensured the sample covered the probe tip and acquired a stable reading.

### Blood pressure measurement

BP was measured using radiotelemetry (probe TA11PA-C10, Data Sciences International, using the Ponemah software, Data Sciences International) and tail-cuff in a CODA non-invasive blood pressure system (Kent Scientific Corporation), as previously decribed.^10,34^

### CD8^+^ T cells adoptive transfer

To isolate CD8^+^ T cells, single-cell suspensions of splenocytes were subject to Dynabeads™ Untouched™ Mouse CD8 Cells Kit (Thermo Fisher Scientific) for the isolation of CD8^+^ leukocytes according to the manufacturer’s manuals. 5×10^6^ of either WT or *Gpr65^−/−^* CD8^+^ T cells were resuspended in 200μL PBS and intravenously injected into *Rag1^−/−^*mice.

### Angiotensin-II induced hypertension model

Six-week-old male WT and *Gpr65^−/−^* mice fed low or high-fiber diets received a subcutaneous minipump (Alzet model 2004) containing angiotensin II (Ang II, 0.5 mg/kg body weight/day; Auspep) placed during surgery under anesthesia with isoflurane. Data were collected for 4 weeks after the implant. Littermate mice were randomized into either group using an Excel randomization tool.

For the adoptive transfer model, one day after the adoptive transfer of the CD8^+^ T cells, ten-week-old male *Rag1^−/−^* recipient mice underwent a subcutaneous minipump (Alzet model 1002) implantation containing Ang II at 1.44 mg/kg body weight/day (Auspep) for 2 weeks as described above. Mice were randomized into either group using an Excel randomization tool.

### Metabolic characterization

Mice were individually housed in metabolic cages for 24 hours. Body weight, food intake, water intake, urine excretion and feces excretion were measured. Mice body composition was analyzed by an EchoMRI 3-in-1 machine (Houston, TX).

### Saline challenge studies

To examine if GPR65 deficiency affected diuretic and natriuretic responses, we performed a saline challenge test according to the protocol kindly shared by Prof. Alicia McDonough (Keck School of Medicine of the University of Southern California).^35,36^

### Cardiac ultrasound

The day before the endpoint, mice were anesthetized using isoflurane, and B- and M-mode echocardiography was performed to image the left ventricle using a Vevo 2100 Ultra High Frequency ultrasound system and an MS550D transducer by an experienced user at the Monash Biomedical Imaging Centre (E.S.).

### Histological analysis

Mouse heart, kidney, and gut tissues were fixed in 10% neutral buffered formalin for 24-48 hours, processed, paraffin-embedded and sectioned at 4 μm. Masson’s trichrome staining was performed to analyze the collagen present in heart, kidney, and intestinal samples. Periodic acid-Schiff alcian blue (PAS/AB) staining was performed to analyze goblet cells in the gut.

### Flow cytometry

Single cell suspensions were isolated from mouse spleen, kidney, and colon (See Online Supplemental files for details). Cells were resuspended with FACS buffer (PBS containing 2% FCS and 4 mM EDTA). Cell viability was evaluated using the fixable viability stain 620 (BD Horizon). After blocking Fc receptors with mouse FcR blocking reagents (Miltenyi Biotech) for 15 mins at room temperature, single-cell suspensions were surface-stained at 4°C for 30 min. Cells were then fixed and permeabilized at 4°C using a Foxp3/Transcription Factor Staining Buffer Set (eBioscience). Intracellular staining was then performed. Cell counts were determined using flow cytometry counting beads (CountBright Absolute; Life Technologies) following manufacturer instructions. Flow cytometry was performed to quantify tissue-specific immune cells and assess pCREB levels and cytokine production (See Online Supplemental Files for details). Sample data were acquired using a five-laser BD LSRFortessa X-20 flow cytometer and BD FACSDiva software (BD Biosciences) and analyzed using FlowJo software (Tree Star).

### *In vitro* cell stimulation

Mouse splenic or colon lamina propria cells were cultured in RPMI1640 medium supplemented with 10% FBS and 4 mM L-glutamine, which was altered to pH 6.5 or pH 7.5 using HCl and NaOH. Cells were stimulated with 100 ng/ml PMA (Sigma-Aldrich) and 1 mg/ml ionomycin (Sigma-Aldrich) for 2-h for pCREB detection, and for 4-h for intracellular cytokine analyses.

### cAMP assay

1X10^6^ splenic cells were resuspended in 200 μL RPMI1640 supplemented with 10% FBS and 4 mM L-glutamine at pH 6.5 or pH 7.5. After incubating at 37°C for 30 mins, cells were washed with PBS at pH 6.5 or pH 7.5, respectively. Intracellular cAMP levels were measured by Invitrogen cAMP Competitive ELISA kit (ThermoFisher, #Cat: EMSCAMPL), following the manufacturer’s instructions.

### DNA extraction from cecal contents and feces

DNA extraction of cecal/fecal contents was performed using the DNeasy PowerSoil DNA isolation kit (Qiagen, Germany). DNA samples were quantified using Nanodrop (Thermo Fisher Scientific).

### Mouse fecal bacterial load measurement

Absolute bacterial load was measured using quantitative real-time PCR. Briefly, we used Fast SYBRgreen (Thermo Scientific) and 16S primers 1114F/1221R in a QuantStudio 7 qPCR Instrument (Thermo Scientific). Samples were loaded at 10ng/well, amplified in triplicates and compared to a standard containing 10^2^ to 10^10^ bacterial copies.

### Short-chain fatty acids quantification by liquid chromatography-mass spectrometry (LC-MS)

Quantification of SCFAs including acetate, propionate, butyrate, isobutyrate, isovalerate, valerate and caproate was performed using LC-MS method, as previously described (See Online Supplemental Files for details).^37^

### Animal statistical analyses

All graphical representation of data was done using Prism 8.2.1 (GraphPad Software, La Jolla, CA). The normality of the distribution of continuous data was assessed by the Shapiro-Wilk test. Comparison between multiple groups of different treatments and different mouse strains was analyzed using a two-way analysis of variance (not for repetitive measures for 4 groups or for repetitive measures for weekly BP) with adjustment using Tukey’s multiple comparisons test. Data is presented as mean ± standard error of the mean*. P*<0.05 was considered significant. All data was collected and analyzed blindly, where possible.

### 16S ribosomal RNA sequencing

V4 region of the 16S ribosomal RNA (rRNA) was targeted and amplified using primers previously published.^38^ 240ng of the amplicon product per sample were pooled and purified. Sequencing was performed at the Australian Genomic Research Facility (AGRF, Melbourne, Australia) on MiSeq instrument (Illumina) to generate 300bp paired-end reads. Sequencing followed our protocol published elsewhere.^39^

## Results

### Dietary fiber lowers large intestinal pH, and this is dependent on the gut microbiota

Mice intestinal pH profiles have mostly been reported under normal conditions without dietary interventions^32^ and have not been well characterized, particularly for the large intestine. To determine the impact of dietary fiber on the large intestinal luminal and interstitial tissue pH, we fed WT mice a control diet and diets high and low in fermentable fiber for 7 days (Figure 1A). Besides dietary fiber, the nutritional components of these diets are almost identical (Table S2). The cecum and colon luminal pH was ∼7.2 when mice were fed the control diet (Figure 1B, Figure S1A). Notably, the cecal and colonic pH were reduced by high fiber diet to 6 and 6.5, respectively (Figure 1B). This change in pH was mirrored by changes in colonic interstitial fluid pH (Figure 1B). This was further accompanied by a significant increase in the acetate and total SCFA levels in the colonic contents (Figure 1C). On the contrary, relative to a high fiber diet, a low fiber diet increased the luminal pH in the colon to ∼7.5 (Figure 1B) and reduced the levels of SCFAs in both the colonic contents and portal vein plasma (Figure 1C and 1D). However, these short-term dietary interventions (7 days) did not alter the levels of SCFAs in the peripheral circulation (Figure S1B), suggesting it is the colon where fiber intake produces marked effects. Aligned with previous reports,^5,11,40^ these dietary interventions significantly altered the composition of the gut microbiome in mice, as indicated by both β-diversity (Figure S1C) and α-diversity (Figure S1D-G), with SCFA producers, such as Lachnospiraceae-NK4A136-group, being enriched with the high fiber diet (Figure S1H-I). Low fiber diet significantly enriched *Alistipes* spp. (Figure S1I), which is associated with intestinal inflammation and hypertension.^41,42^

**Figure. 1.**
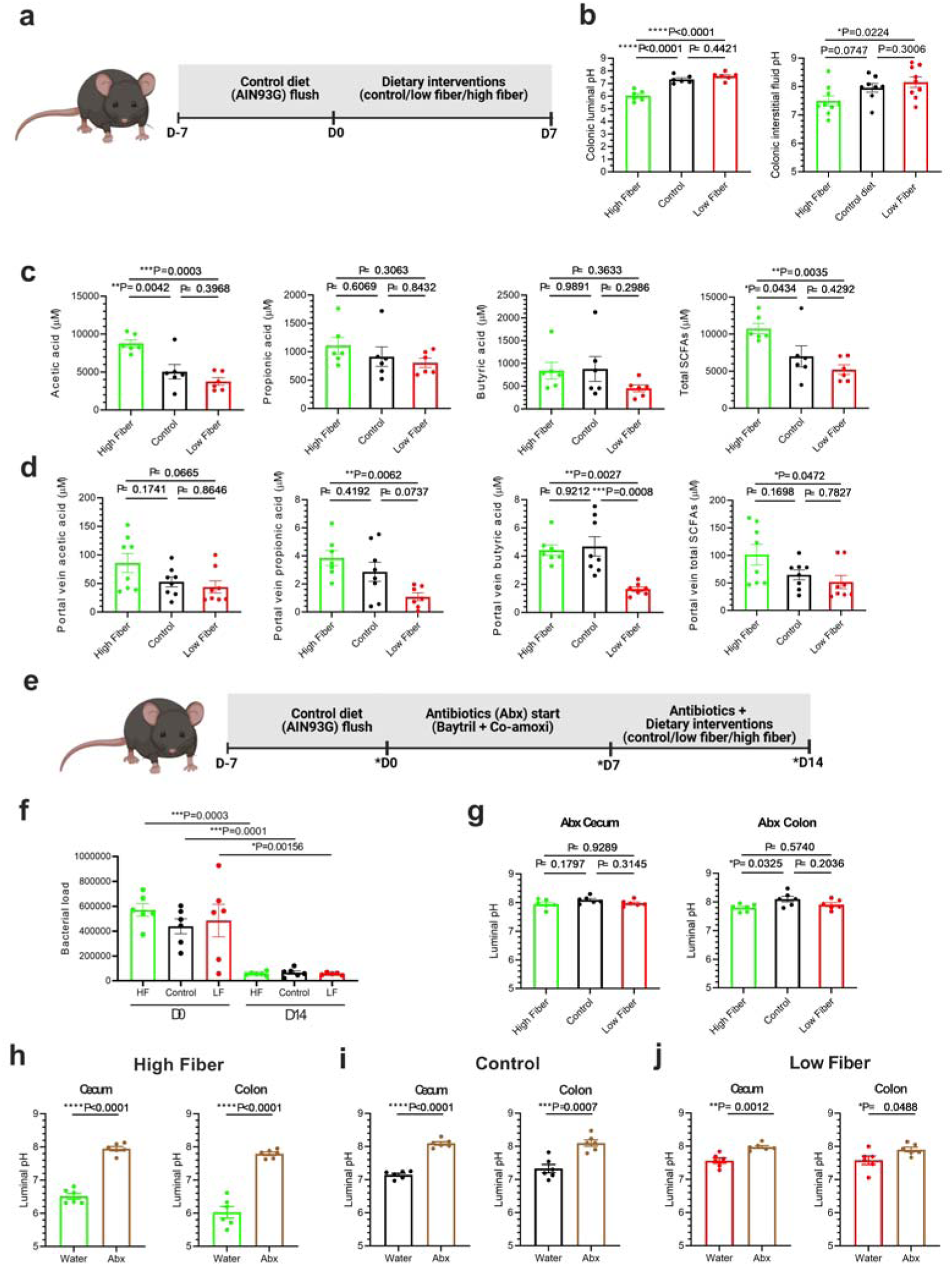
Dietary fiber lowers colonic pH through the gut microbiota. **A**, The experiment design to characterize the impact of dietary fiber on luminal pH in cecum and colon. **B**, Colonic luminal and interstitial fluid pH at the endpoint, post dietary interventions. **C**, Levels of acetic, propionic, butyric acids and total short-chain fatty acids (SCFAs) in cecal contents and **D**, in plasma from the portal vein. **E**, The experiment design to characterize the impact of dietary fiber on luminal pH in the cecum and colon with antibiotic treatment. Time points with asterisks indicate when fecal samples were collected for bacterial load measurements. **F**, Bacterial load. **G**, Cecal and colonic luminal pH at the endpoint, post dietary interventions with antibiotics treatments. **H**, Each data point represents an individual mouse, and n=6 mice in each group for all panels besides colonic interstitial fluid pH (n=8 for control diet, n=9 low and high fiber diets); all data represented as means ± SEM; **C**, **D**, **G**, One-way ANOVA with adjustment for multiple comparisons; **H**, Student’s t test; *P<0.05, **P<0.01, ***P<0.001, ****P<0.0001.

We then aimed to determine whether the pH-lowering effect of dietary fiber is dependent on the gut microbiota. To achieve this, we pre-treated mice with an antibiotic cocktail to reduce the bacterial load before starting the same dietary interventions (Figure 1E). Indeed, the antibiotic cocktail effectively reduced the bacterial load (Figure 1F). This reduction in bacteria in the intestine resulted in a higher luminal pH in the cecum and colon, independent of the level of fiber in the diet (Figure 1G-J). This is consistent with previously reported data showing lacking gut microbiota leads to large intestinal alkalization.^32^ Our data further support the theory that acidic large intestinal pH produced when fiber is consumed is dependent on the gut microbiota, particularly bacteria.

### Acidic pH regulates GPR65 signaling and inhibits pro-inflammatory pathways in both colon and spleen derived CD8^+^ T cells

The colonic luminal pH produced by high fiber intake, pH 6.5, activates a family of proton-sensing GPCRs, while the pH produced by low fiber intake, pH 7.5, silences them.^14,15^{Howard, 2024 #6584} Within this family, we chose to focus on GPR65 due to its previously known association with inflammatory diseases related to gut homeostasis,^18–23^ as well as its more recent association with hypertension in PheWAS studies (Table S1).^22^ GPR65 expression is particularly enriched in the immune compartment.^27–29^ There are no robust antibodies against GPR65, so we leveraged a mouse model where GFP replaced one copy of the GPR65 gene to detect where GPR65 was being expressed. Using *Gpr65^gfp/+^* mice,^22,43^ we found that CD8^+^ T cells highly expressed GPR65 protein compared to CD4^+^ T cells (Figure 2A). This is consistent with human GPR65 expression in a public database.^28^ Interestingly, CD8^+^ T cells causally contribute to the elevation of BP.^36^ Moreover, CD8^+^ T cells extracted from colon lamina propria had a higher level of GPR65 expression compared to splenic CD8^+^ T cells, while CD4^+^ T cells extracted from colon lamina propria and spleen did not differ in GPR65 expression (Figure 2A). In addition, we found that *Gpr65^−/−^* mice had significantly larger spleens (Figure 2B) with significantly more leukocytes, including CD4^+^, CD8^+^ and γδ T cells (Figure 2C, Figure S2A and S2B), but not Foxp3^+^ regulatory T cells (T_reg_) (Figure S2C).

**Figure 2.**
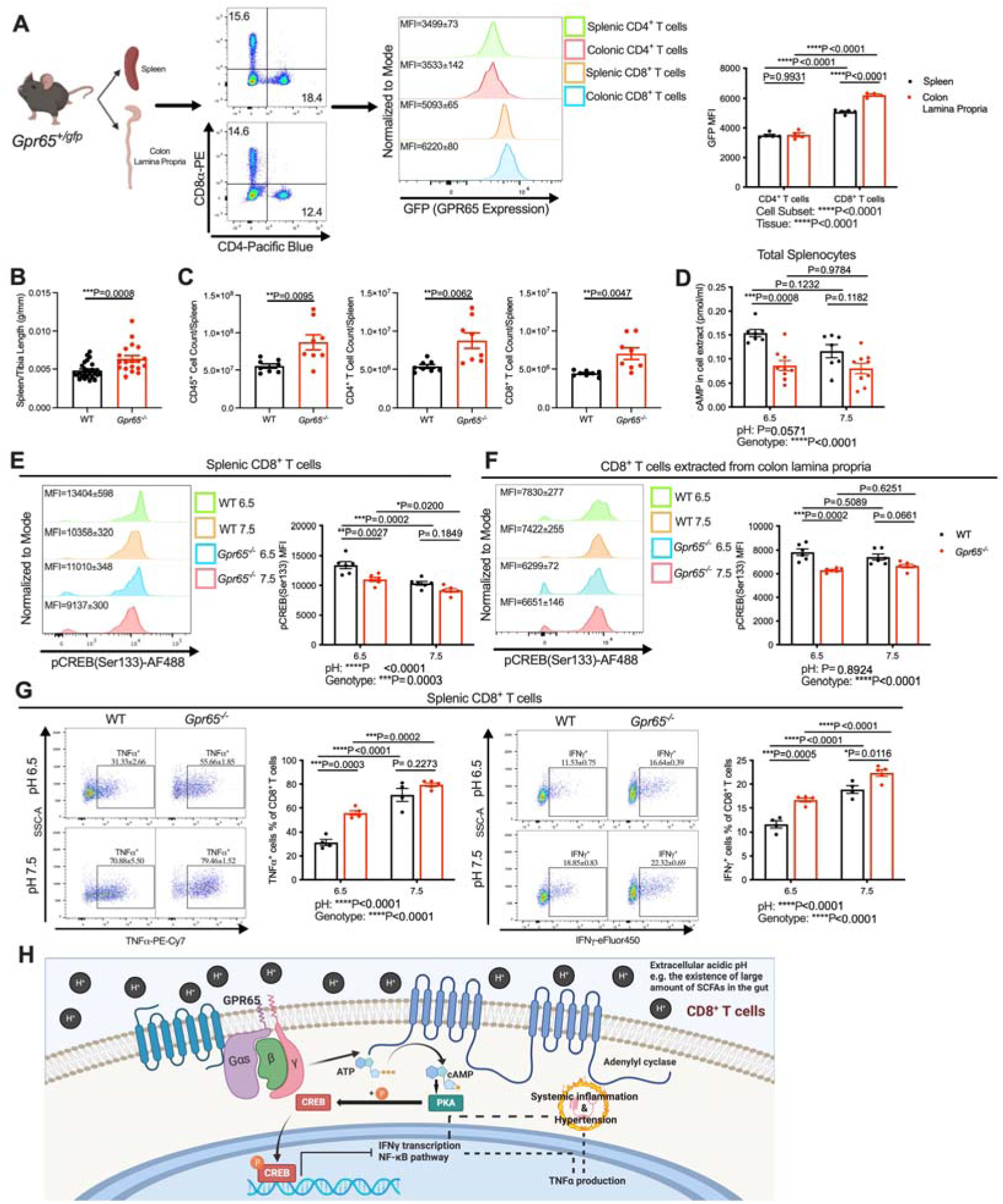
pH and GPR65 signaling modulate immune responses related to hypertension pathogenesis. Single cell suspensions were prepared from the spleens or colon lamina propria of male *Gpr65^gfp/+^* mice at the age of 12 weeks. **A**, GPR65 expression in CD4^+^ and CD8^+^ T cells from spleen and colon lamina propria indicated as mean fluorescence intensity (MFI) of GFP. Single cell suspensions were prepared from the colon lamina propria of male wild-type C57BL/6J (WT) mice or *Gpr65^−/−^*mice at the age of 12 weeks. **B**, CD4^+^ and CD8^+^ T cell count in the colon lamina propria. Colon lamina propria cells were incubated with PMA and ionomycin for 2 hours. **C**, pCREB (Ser133) levels of CD8^+^ T cells extracted from colon lamina propria. Colon lamina propria cells were incubated with PMA and ionomycin for 4 hours. **D**, Percentage of TNFα producing CD8^+^ T cells at pH 6.5 and 7.5 among WT and *Gpr65^−/−^* CD8^+^ T cells derived from colon lamina propria. **E**, Percentage of IFNγ producing CD8^+^ T cells at pH 6.5 and 7.5 among WT and *Gpr65^−/−^* CD8^+^ T cells derived from colon lamina propria. **B**, Spleen weight to tibia length index from male mice at the age of 12 weeks. **C**, CD45^+^ immune cell count, CD4^+^ and CD8^+^ T cell count in the spleen. Splenocytes were isolated from mice at the age of 8-10 weeks and incubated under either pH 6.5 or pH 7.5 for 30 min. **D**, Levels of cAMP in the cell lysate extracts. **E**, pCREB (Ser133) levels of splenic CD8^+^ T cells in splenocytes incubated with PMA and ionomycin for 2 hours. **F**, pCREB (Ser133) levels of CD8^+^ T cells extracted from colon lamina propria incubated with PMA and ionomycin for 2 hours. **G**, Percentage of TNFα and IFNγ producing CD8^+^ T cells at pH 6.5 and 7.5 among WT and *Gpr65^−/−^* splenocytes were incubated with PMA and ionomycin for 4 hours. **H**, Graphic summary of how pH regulates inflammatory responses contributing to hypertension in CD8^+^ T cells. Created with BioRender.com. Each data point represents an individual mouse, and n=4-26 mice in each group; All data represented as means ± SEM. **A**, **D-G**, Two-way ANOVA with adjustment for multiple comparisons. **B**, **C**, Student’s t test. *P<0.05, **P<0.01, ***P<0.001, ***P<0.0001.

GPR65 is a Gαs-coupled receptor whose activation induces intracellular cAMP accumulation.^44^ cAMP production in response to low pH was impaired in macrophages of GPR65 deficient mice.^45^ We discovered that low (GPR65 activating) pH (6.5) increased the level of intracellular cAMP in splenic cells in a GPR65-dependent manner (Figure 2D). The increased intracellular cAMP can lead to the phosphorylation of cAMP response element-binding protein (CREB). Phosphorylated CREB (pCREB) has various roles in immune responses, including suppressing the production of pro-inflammatory cytokines, tumor necrosis factor alpha (TNFα) and interferon gamma (IFNγ),^46^ which underlie the pathogenesis of hypertension.^47,48^ Thus, we investigated whether pH and GPR65 signaling regulated the phosphorylation of CREB in T cells extracted from the spleen and colon lamina propria *in vitro*. Compared to pH 7.5, pH 6.5 significantly elevated pCREB levels in WT splenic CD8^+^ and CD4^+^ T cells activated by PMA and ionomycin (Figure 2E and Figure S2D). The elevation of pCREB by acidic pH was impaired in *Gpr65^−/−^* splenic CD8^+^ T cells (Figure 2E), but not splenic CD4^+^ T cells (Figure S2D). While we did not observe a significant difference in pCREB levels in WT colonic T cells activated by PMA and ionomycin at pH 6.5 and pH 7.5, GPR65 deficiency significantly reduced the pCREB levels of colonic CD4^+^ and CD8^+^ T cells under pH 6.5 but not pH 7.5 (Figure 2F and Figure S2E). Downstream of the GPR65-pCREB axis, acidic pH 6.5 indeed suppressed TNFα and IFNγ production from splenic and colonic CD8^+^ T cells in a GPR65-dependent manner (Figure 2G and S2F). However, the impact of acidic pH and GPR65 signaling was less profound on TNFα and IFNγ production from splenic and colonic CD4^+^ T cells (Figure S2G and S2H).

In summary, pH change, which resembles the regulation of colonic luminal pH by dietary fiber, regulates GPR65 signaling. Low pH, via GPR65 activation, suppresses the pro-inflammatory cytokine production by CD8^+^ T cells both locally from the colon and systemically from the spleen (Figure 2H). We hypothesized this may reduce BP and protect against CVD.

### Deficiency of GPR65 does not impair gut homeostasis

We then examined the impact of GPR65 deficiency on gut homeostasis. *Gpr65^−/−^* and WT mice exhibited similar gut microbiome compositions (Figure S3A and S3B), and immune cell composition in the colon, including CD4^+^, CD8^+^, γδ T cells, and T_reg_ cells (Figure S3C and S3D). However, *Gpr65^−/−^* mice had altered gut physiology (Figure S3E), demonstrated in fibrosis (Figure S3F), muscularis propria thickness (Figure S3G) and goblet cell frequency (Figure S3H) compared to WT mice.

### Deficiency of GPR65 results in spontaneous cardiovascular dysfunction

The role of GPR65 in the cardiovascular system and BP regulation remains largely unknown. Therefore, we characterized multiple key cardiovascular phenotypes of *Gpr65^−/−^* mice. Compared to WT mice, male *Gpr65^−/−^* mice had no difference in body weight (Figure S4A), body composition (Figure S4B through S4D), food and water intake (Figure S4E and S4F), or feces and urine excretion (Figure S4G and S4H). We discovered, however, that *Gpr65^−/−^* mice spontaneously developed higher diastolic BP compared to WT mice (Figure 3A). Consistently, *Gpr65^−/−^* mice had significantly enlarged heart and kidney compared to WT mice (Figure 3B and 3C). This was accompanied by significantly higher levels of cardiac fibrosis (Figure 3D and 3E), left ventricular posterior wall in the end diastole (LVPW;d, Figure 3F), and cardiac output (Figure 3G). Similarly, renal fibrosis was significantly elevated in *Gpr65^−/−^* mice (Figure 3H and 3I). We performed a saline challenge to investigate the influence of GPR65 deficiency on renal function. Compared to WT mice, *Gpr65^−/−^* mice excreted less water and sodium over a 5-hour period (Figure 3J and 3K), which suggests impaired diuretic and natriuretic functions. Aligned with the significant functional phenotypes, urine creatinine levels were significantly increased in *Gpr65^−/−^* mice (Figure 3L). We then investigated the number of immune cells in the kidney (Figure S4I and S4J). We observed significantly elevated CD45^+^ immune cells (Figure 3M), particularly CD8^+^ T cells (Figure 3N). CD4^+^ T cells (Figure 3O), γδ T cells (Figure 3P), macrophages/monocytes (Figure 3Q), and dendritic cells (Figure S4K) were also more abundant in the kidneys of *Gpr65^−/−^* mice, while no difference in T_reg_ cells (Figure S4L) and neutrophils (Figure S4M) were observed.

**Figure 3.**
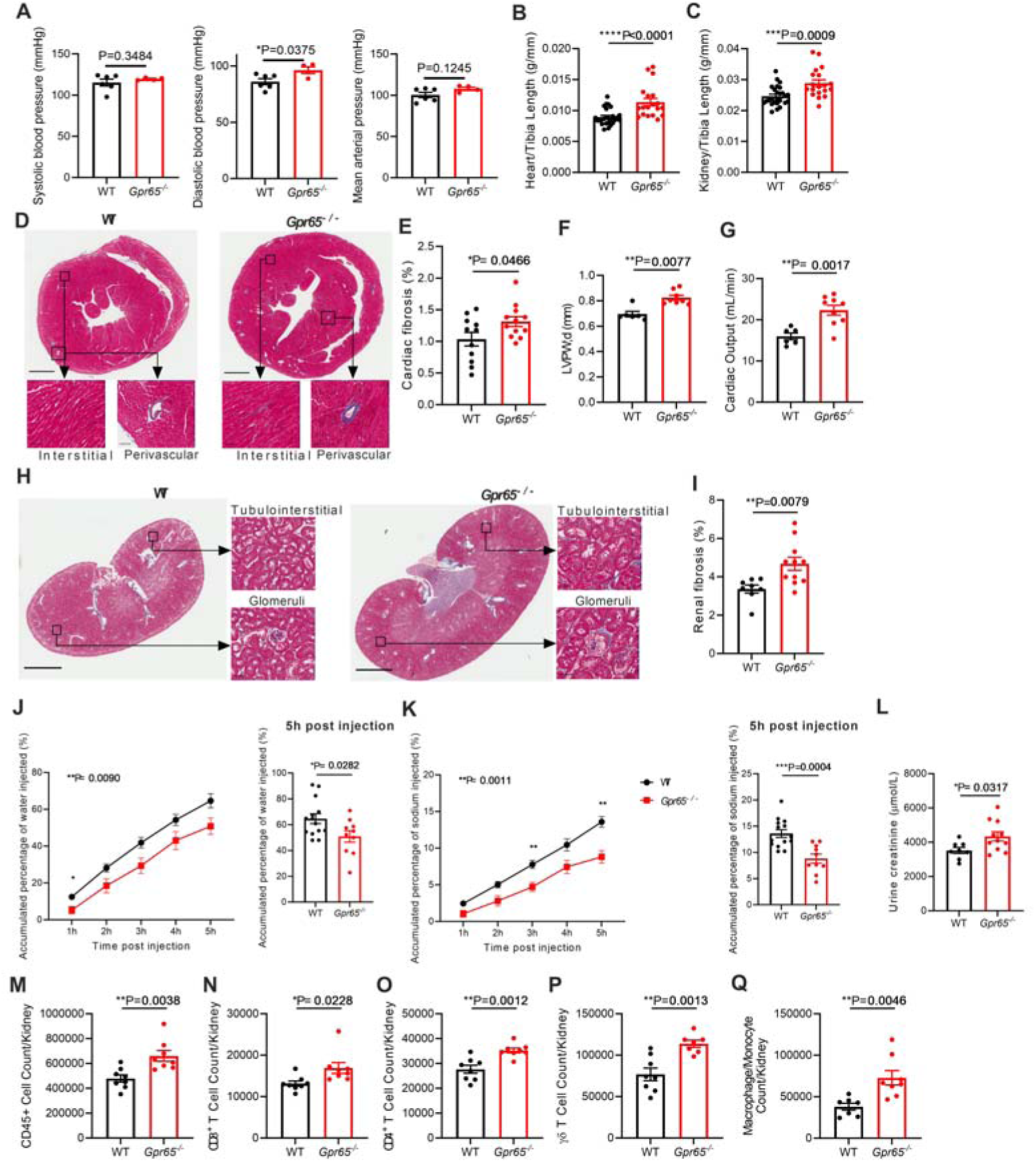
GPR65 deficiency spontaneously increases blood pressure, tissue hypertrophy, and renal inflammation in male mice. **A**, Systolic blood pressure (BP) of male mice measured non-invasively by radio-telemetry at the age of 10 weeks. Tissues of male mice were collected and weighed at the age of 12 weeks. **B**, Heart weight to tibia length index. **C**, Kidney weight to tibia length index. **D**, Representative heart sections stained with Masson’s trichrome; For upper panels, magnification = ×10, scale bar = 1mm; For lower left panels (interstitial), magnification = ×200, scale bar = 50μm; For lower right panels (perivascular), magnification = ×100, scale bar = 100μm. **E**, Percentage of fibrotic area in the cardiac tissues. **F**, Left ventricle posterior wall in the end diastole and **G**, cardiac outputs from ultrasound screening was performed at 11 weeks of age. **H**, Representative kidney sections stained with Masson’s trichrome; For left panels, magnification= ×5, scale bar = 2mm; For right panels, magnification = ×200, scale bar = 50μm. **I**, Percentage of fibrotic area in the renal fibrosis. Saline challenge was performed at the age of 11 weeks. **J**, Renal diuretic functions measured by accumulated water excretion over 5 hours post injection. **K**, Renal natriuretic functions measured accumulated sodium excretion over 5 hours post injection. **L**, The level of creatinine in urine from urine were collected from 24h metabolic caging. **M**, CD45^+^ immune cell count, **N**, CD8^+^ T cell count, **O**, CD4^+^ T cell count, **P**, γδ T cell count and **Q** Macrophage/Monocyte count from kidneys of male mice at 12 weeks of age. For the left figures of panels **J** and **K**, n=10-14 mice in each group; All data represented as means ± SEM; Two-way ANOVA. For other panels, each data point represents an individual mouse, and n=6-26 mice in each group; All data represented as means ± SEM; **B, C**, **F**, Mann-Whitney test; All other panels, Student’s t test. *P<0.05, **P<0.01, ***P<0.001, ****P<0.0001.

Importantly, we observed similar cardiovascular phenotypes in female *Gpr65^−/−^* mice, including higher BP (Figure S5A), cardiorenal hypertrophy (Figure S5B and S5C), thicker LVPW;d (Figure S5D), and elevated cardiorenal fibrosis (Figure S4E through S4H) compared to female WT mice. Collectively, these results clearly demonstrated that GPR65 deficiency leads to the spontaneous development of high BP and associated end-organ damage which leads to CVD.

### Deletion of GPR65 decreased the protection of dietary fiber against high blood pressure

We next investigated whether pH-sensing by GPR65 conferred cardiovascular protection by dietary fiber. To achieve this, we challenged WT and *Gpr65^−/−^* mice with Ang II (0.5mg/kg body/weight/day) and fed the mice either high or low fiber diets (Figure 4A). As previously reported by us, high fiber diet suppressed the elevation of systolic BP (Figure 4B) induced by Ang II infusion in WT mice.^5^ Such suppression was significantly smaller in *Gpr65^−/−^* mice (Figure 4B). GPR65 deficiency significantly increased BP with hypertensive stimuli when mice were fed with high fiber diet, while the impact of GPR65 genotype was negligible under low fiber diet (Figure 4B). GPR65 deficient mice fed high fiber had higher cardiac weight (Figure 4C), thicker LVPW;d (Figure 4D), reduced ejection fraction (Figure 4E) and fractional shortening (Figure 4F), and higher cardiac fibrosis relative to WT mice (Figure 4G and 4H). Similarly, kidney weight (Figure 4I) and renal fibrosis (Figure S6A and S6B) were increased. Renal natriuretic and diuretic function were also impaired by the GPR65 deficiency (Figure 4J and 4K). We quantified the number of immune cells in the kidneys and observed that high fiber fed *Gpr65^−/−^* mice had significantly more CD45^+^ immune cells than the high fiber fed WT mice (Figure S6C). Notably, the infiltration of CD8^+^ T cells, whose responses were regulated by acidic pH and GPR65 signaling, was significantly higher in the kidney of *Gpr65^−/−^* mice (Figure 4L). In addition, γδ T cells and macrophages/monocytes were also increased (Figure S6D and S6E), while CD4^+^ T cells, T_reg_ cells, neutrophils and dendritic cells were not (Figure S6F through S6I). However, when mice were fed with a low fiber diet, which mimics an environment where GPR65 is not activated, there was no differences in cardiac or renal parameters besides fibrosis, which was further exacerbated in *Gpr65^−/−^* mice (Figure 4B through 4K, Figure S6A and S6B). Consistent with this, no difference was observed in immune cell infiltration in the kidney under a low fiber diet (Figure 4L, Figure S6C through S6I).

**Figure 4.**
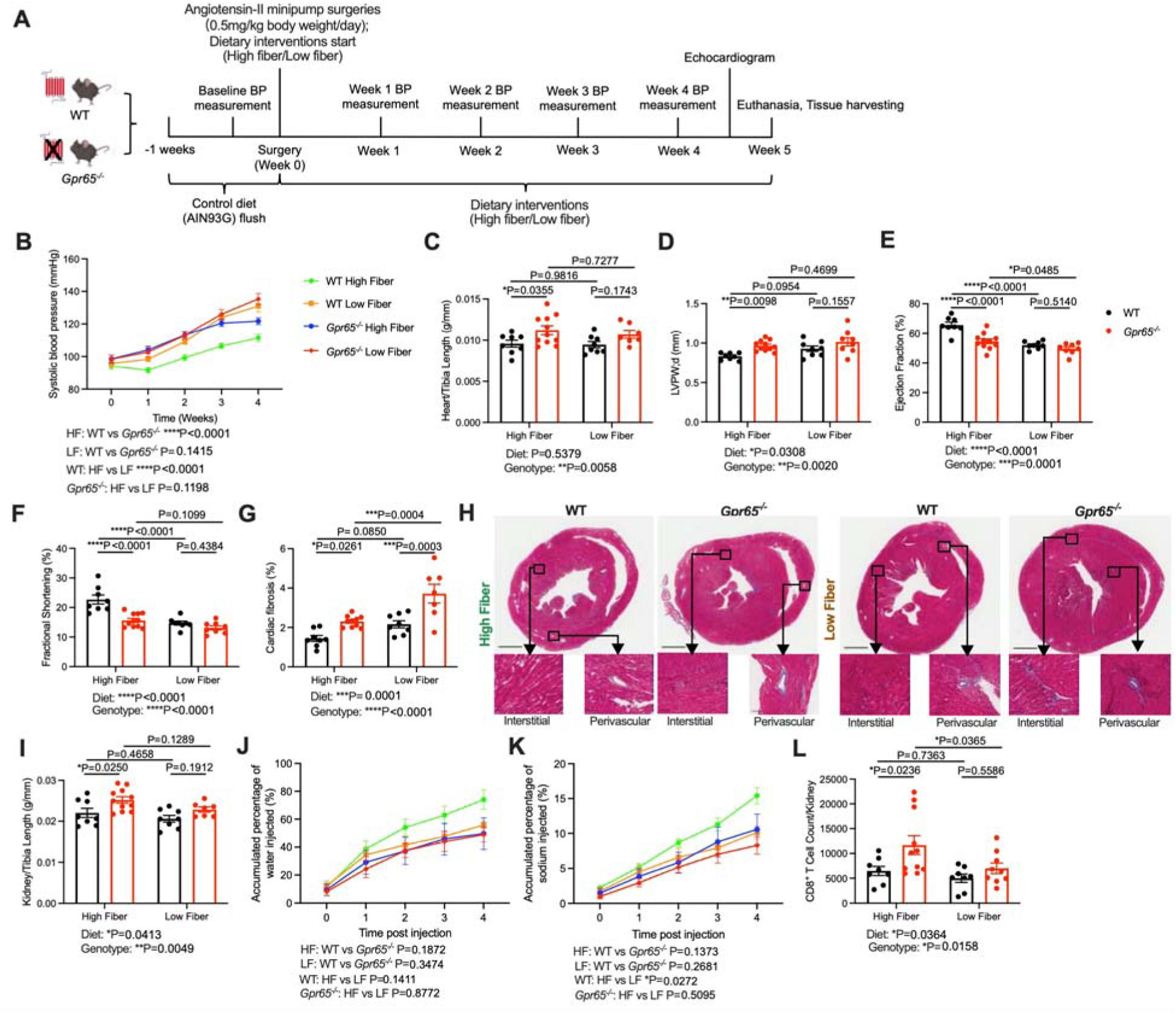
Deficiency of GPR65 decreased the cardiorenal protective effects of dietary fiber. 6-week-old male WT and *Gpr65^−/−^* were infused with 0.5 mg/kg body weight/day angiotensin-II for 28 days and fed with either high fiber (HF) or low fiber (LF) diet. **A**, The experiment design. **B**, Systolic BP profile over 4 weeks of infusion; **C**, Heart weight to tibia length index; Cardiac ultrasound scanning was performed at week 4 of infusion. **D**, Left ventricle posterior wall in the diastole; **E**, Cardiac ejection fraction; **F**, Cardiac fractional shortening; **G**, Percentage of fibrotic area in the cardiac tissues stained with Masson’s trichrome. **H**, Representative heart sections; For upper panels, magnification = ×10, scale bar = 1mm; For lower left panels (interstitial), magnification = ×200, scale bar = 50μm; For lower right panels (perivascular), magnification = ×100, scale bar = 100μm. **I**, Kidney weight to tibia length index. Mice were intraperitoneally injected with 10% of their body weight of 37 °C 0.9% saline solution at the 3^rd^ week post minipump implantation. **J**, Renal diuretic and **K,** natriuretic functions measured by accumulated water excretion over 5 hours post injection. **L**, CD8^+^ T cell count from kidneys of the mice at the endpoint of the study. Each data point represents an individual mouse, and n=8-11 mice in each group; All data represented as means ± SEM. Two-way ANOVA with adjustment for multiple comparisons. *P<0.05, **P<0.01, ***P<0.001, ****P<0.0001.

We then examined the large intestinal tissue of mice fed low and high fiber diets challenged with Ang II. High fiber intake thickened the muscularis propria layer and increased the length of intestinal villi in the cecum, irrespective of *Gpr65* genotype (Figure S7A though S7C) but did not influence the width of the fibrotic layer and the number of goblet cells (Figure S7D and S7E). We also characterized the gut microbiome composition by analyzing the 16S rRNA gene. The main factor driving the differences in gut microbiota was diet rather than the genotype for *Gpr65* (Figure S7F and S7G). The top leading taxonomic change explaining the differences in gut microbiome was the enrichment of *Bacteroides* in the groups fed high fiber diet (Figure S7H and S7I). *Bacteroides* species are the most predominant anaerobes in the gut^49^ with significant cardiovascular benefits.^50–53^ Interestingly, *Bacteroides* spp. are the dominant carbohydrate fermenters at pH 6.0 to 7.0, the normal luminal pH within the cecum and colon.^54–56^ The major products of carbohydrate fermentation by *Bacteroides* spp. are propionate and acetate, rather than butyrate.^57,58^ Indeed, compared to low fiber, high fiber diet profoundly elevated acetate and propionate levels in the cecal contents, independent of the genotype of the mice, while no significant difference in cecal butyrate levels was observed (Figure S7J). *Lachnospiraceae*-NK4A136-group, enriched following the 7-day high fiber intervention (Figure S1D), was also more abundant in mice fed high fiber diet in this study (Figure S7I). Low fiber diet expanded the genera *Bilophila* and *Alistipes* (Figure S7H and S7I), which are associated with inflammation and higher risk of CVD.^41,42,59,60^

### GPR65 absence in CD8^+^ T cells elevated blood pressure under high fiber diet

We then investigated whether immune regulation by acidic pH and GPR65 in CD8+ T cells causally contributed to elevated BP under a high-fiber diet. *Rag1^−/−^* mice lack both T and B cells and have blunted BP elevation in the Ang II-induced hypertension model.^61^ We adoptively transferred splenic CD8^+^ T cells from WT or *Gpr65^−/−^* mice fed a high fiber diet (Figure S8A) into *Rag1^−/−^* mice which were also fed high fiber diet, then infused with Ang II for 14 days (Figure 5A). We indeed observed that *Rag1*^−/−^ mice that received *Gpr65^−/−^* CD8^+^ T cells developed higher BP than *Rag1^−/−^* mice that received WT CD8^+^ T cells (Figure 5B). *Gpr65^−/−^* CD8^+^ T cells also led to cardiac hypertrophy in Ang II-infused *Rag1^−/−^* mice but had no significant impact on kidney size (Figure 5C). Using flow cytometry, we confirmed the presence of adoptive transferred CD8^+^ T cells in the recipient mice’s kidneys, blood, and colon lamina propria (Figure S8A). We also observed a trend towards more abundant CD8^+^ T cells in the kidneys of *Rag1^−/−^* mice that received *Gpr65^−/−^* CD8^+^ T cells (Figure 5D) but not in blood and colon lamina propria (Figure S8B and S8C). Moreover, *Gpr65^−/−^* CD8^+^ T cells resulted in significantly higher CD45^+^ leukocyte infiltration, particularly macrophage/monocytes (Figure 5E). We did not observe these changes in blood and colon lamina propria (Figure S8D and S8E). Together, these results demonstrated that GPR65 signaling in CD8^+^ T cells causally regulates BP and improves the accompanied cardiovascular complications under a high fiber diet.

**Figure 5.**
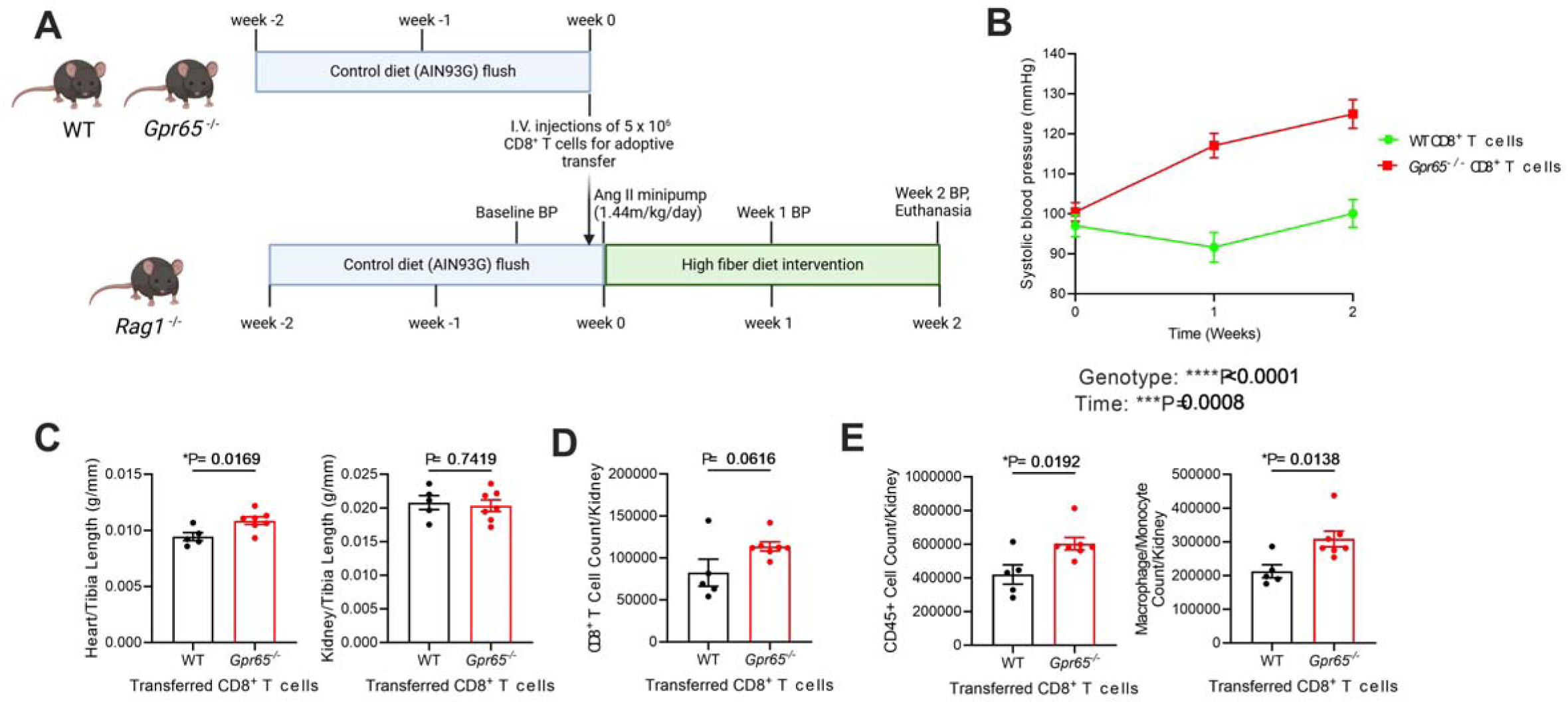
Deficiency of GPR65 in CD8^+^ T cells causes BP elevation in mice under the high fiber diet. 10-week-old male *Rag1^−/−^* mice were adoptively transferred with either WT and *Gpr65^−/−^* CD8^+^ T cells, and then infused with 1.44 mg/kg body weight/day angiotensin-II for 14 days. They were fed with HF diet. **A**, The experimental design; **B**, Systolic BP profile over 2 weeks of infusion; **C**, Heart weight to tibia length and kidney weight to tibia length indexes. **D**, CD8^+^ T cell count, **E**, CD45^+^ immune cell count and Macrophage/Monocyte count from kidneys of the mice at the endpoint of the study. Each data point represents an individual mouse, and n=5-7 mice in each group; All data represented as means ± SEM. **B**, Two-way ANOVA with adjustment for multiple comparisons. All other panels, Student’s t test. *P<0.05, ***P<0.001, ****P<0.0001.

## Discussion

Dietary fiber and SCFAs are potent BP-lowering agents. Promoting dietary fiber or SCFAs intake represents a novel, safe, and affordable therapy against hypertension. SCFAs, particularly acetate, propionate, and butyrate, are well-known for their immunomodulatory effects by activating metabolite-sensing GPCRs, such as GPR41, GPR43 and GPR109A.^5,9,10^ Some evidence suggests these receptors may confer cardiovascular protection.^5,9,10^ Here, we identified a novel mechanism of how dietary fiber regulates BP and cardiovascular health via colonic luminal pH and GPR65. We show that a high-fiber diet lowers large intestinal pH by increasing the production of SCFAs by bacteria. This reduction in large intestinal pH activates the proton-sensing receptor GPR65, particularly in CD8^+^ T cells, and its downstream anti-inflammatory signaling cascade, which lowered BP elevation induced by Ang II. Together, our findings support that large intestinal pH-sensing by GPR65, at least partly, explains the systemic cardiovascular protection by dietary fiber.

Colonic epithelial cells are naturally acidic. This largely results from the microbial production of acidic metabolites, particularly SCFAs.^11–13^ Acidic pH in the colon is essential in maintaining gut homeostasis by suppressing the growth of pathogens and facilitating the expansion of beneficial microorganisms, including SCFA producers.^62^ Indeed, we used dietary interventions to show that fiber determined the luminal pH in the colon and that it was inversely associated with the levels of SCFAs in both the large intestine and portal vein plasma. Such regulation was dependent on the presence of bacteria in the colon.

Among the host’s pH sensors, GPR65 has critical roles in gastrointestinal homeostasis.^18–24,63^ GPR65 is particularly enriched in immune cells and lymphoid tissues.^27–29^ Evidence built in the last decade strongly supports the contribution of inflammation in the development of hypertension,^47,48^ making hypertension an unconventional inflammatory disease. CD8^+^ T cells are a well-known immune cell subset contributing to the pathogenesis of hypertension.^36^ We found GPR65 is more highly expressed by CD8^+^ T cells than CD4^+^ T cells, and the expression was particularly enriched in colon lamina propria. This expression profile suggests this receptor may have a more significant role in CD8^+^ T cells than CD4^+^ T cells, particularly in the colon. Furthermore, our study suggested GPR65 deficiency increased immune cell counts in both the spleen and kidneys. Our data demonstrated that low pH, similar to the large intestinal pH post-high-fiber intervention, suppresses pro-inflammatory cytokine production by CD8+ T cells extracted from the spleen and colon lamina propria via activating GPR65 signaling. This is a plausible mechanism linking dietary fiber, low intestinal pH in the large intestine, GPR65 and cardiovascular protection.

Characterization of its cardiovascular phenotype demonstrated that lack of GPR65 led to the spontaneous development of mild but significant cardiovascular disorders featured by impaired cardiorenal function, and cardiorenal remodeling. These results demonstrate GPR65 has a protective role for cardiovascular health. This phenotype was exacerbated when mice lacking GPR65 were challenged with Ang II, particularly in a low-fibre diet. This shows a novel gene-by-environment interaction. We propose that pH-sensing by GPR65 mediates cardiovascular protection through dietary fiber via anti-inflammatory mechanisms. Our study demonstrates that the cardiovascular protection conferred by dietary fiber is, at least in part, mediated by GPR65. When challenged with Ang II, *Gpr65^−/−^* mice had higher BP and cardiorenal hypertrophy compared to WT controls fed high fiber, and similar BP when fed a low fiber diet, which would also silence GPR65 signaling in the large intestine. Indeed, we observed that high fiber-fed *Gpr65^−/−^* mice had more CD8^+^ T cells in the kidney compared to high fiber fed WT mice. Analysis of the microbiome using 16S rRNA sequencing demonstrated that diet, not *Gpr65* genotype, drove the difference in the gut microbiota composition. High fiber diet promoted *Bacteroides* spp., a genus with known cardiovascular benefits,^50–53^ and increased the levels of acetate and propionate, but not butyrate, in the intestine. Moreover, high fiber also inhibited certain taxa associated with inflammation and cardiovascular disorders, including *Bilophila* and *Aliptipes.*^41,42,59,60^ These data suggested that GPR65 signaling does not regulate gut microbiota composition directly and its effects on cardiovascular health are dependent on downstream pathways triggered by GPR65, such as immunomodulation as discussed above. Indeed, the adoptive transfer model suggested that GPR65 deficiency in CD8^+^ T cells could causally explain the BP elevation under high fiber diet.

The interventions with low and high fiber diets show robust differences in the phenotype between WT and *Gpr65^−/−^* knockout mice, supporting the notion that GPR65 signaling in hypertension starts in the colon. We acknowledge that since we used a systemic knockout model, our animal studies cannot exclude the effect of pH-sensing in other tissues. However, the *in vitro* experiments used the luminal colonic pH we determined *in vivo*; however, we acknowledge these may not correspond to colonic tissue pH.

In conclusion, this is the first study to report the role of colonic luminal and tissue pH on the host’s cardiovascular health. We provided new insights into how dietary fiber and gut microbiota-derived metabolites regulate host health from the perspective of pH. We established a role for GPR65 as a metabolite-sensing GPCR by sensing pH changes caused by microbial fiber fermentation, showing a novel gene-by-environment interaction. Since pH alterations happen universally, this mechanism may extend to broader applications. Agents reducing colonic luminal pH or agonists activating GPR65 may be potential tools for the global control of CVDs.

## Acknowledgements

This study was supported by a National Heart Foundation Vanguard Grant (102927) and a National Health & Medical Research Council of Australia (NHMRC) Ideas Grant (GTN2017382). L.X., M.J.A were supported by Monash Graduate Scholarships. R.R.M was supported by a scholarship from the Faculty of Science, Monash University. J.A.O. is supported by an NHMRC Fellowship (GTN1124288). F.Z.M is supported by a Senior Medical Research Fellowship from the Sylvia and Charles Viertel Charitable Foundation Fellowship, a National Heart Foundation Future Leader Fellowship (105663) and an NHMRC Emerging Leader Fellowship (GNT2017382). The Baker Heart & Diabetes Institute is supported in part by the Victorian Government’s Operational Infrastructure Support Program. The authors thank Monash Animal Research Platform, Monash Genomic Modification Platform, Monash Histology Platform, Monash FlowCore, Monash Biomedical Imaging, Monash Proteomics and Metabolomics Facility, Monash Bioinformatics, and the Australian Genome Research Facility for their technical supports. Summary figures were created with Biorender. We thank Prof Peter Gibson, Jane Muir, Dakota Rhys-Jones, Daniel So, and Chu Yao, for their feedback on the manuscript.

## Author contributions

L.X. planned and performed most of the animal in vivo and in vitro experiments, data analyses, provided intellectual inputs and wrote the manuscript. R.R.M., E.D., S.A., K.C.L., Z.M., K.M.C. contributed to animal in vivo experiments. J.A.O. contributed to pCREB detection. M.P., A.P. performed gut histology analyses. H.J. contributed to animal in vivo experiments and rodent cardiac ultrasound data analyses, heart and kidney histology analyses. E.S. performed rodent cardiac ultrasound scanning and supervised the data analyses performed by L.X.. D.A. performed SCFA measurement by LC-MS. C.A., M.J.A., Y.A.Y. contributed to animal in vivo experiments, mouse strain maintenance and cAMP assay. D.C. supervised SCFA measurement by LC-MS. R.R. contributed to animal in vivo experiments, mouse strain maintenance and provided intellectual inputs. C.R.M. and F.Z.M. conceived the study and supervised the research. F.Z.M. secured the funding to support this study. All authors approved of and contributed to the final version of the manuscript.

