## Supplementary material for "Dietary fiber controls blood pressure and cardiovascular risk by lowering large intestinal pH and activating the proton-sensing receptor GPR65": Online supplemental methods, tables and figures

**Affiliations**

### Methods

#### Phenome-wide association study

A phenome-wide association study (PheWAS) investigated whether *GPR65* was associated with BP and associated end-organ damage by using data available in the Atlas of Genome-Wide Association Study (GWAS) Summary Statistics in 19,516 to 385,699 individuals.[1]

#### Animals

All animal care and experimental procedures used in this study were approved by the Animal Ethics Committee of Monash University (17465, 27929, 37720). Wild-type (WT) C57BL/6J mice were obtained from the Monash Animal Research Platform, Monash University, housed in the same facility under the same conditions. *Gpr65*$^{-/-}$ mice on a C57BL/6J background were generated by using CRISPR/Cas9-based protocol at the Monash Genome Modification Platform (MGMP) at Monash University as previously reported.[2,3] Briefly, the UCSC Genome Browser was used to identify guide RNA target sites flanking the *Gpr65* gene. The following guide RNA were used: 2488 base pairs upstream of the ATG of *Gpr65* (5' GCCTGTTCAAACCCAGCGTG 3') and 401bp downstream of the STOP codon of *Gpr65* (5' CCCGATCACATCCATTATCA 3') to knock out *Gpr65* gene. CRISPR RNAs (crRNA, IDT) were annealed with trans-activating crRNA (tracrRNA) to form a functional crRNA:tracrRNA guide RNA duplex. Cas9 nuclease (IDT, # 1081058) was incubated with the guide RNAs to form a ribonucleoprotein (RNP) complex. Cas9 nuclease (30ng/µl) and the crRNA:tracrRNA guide RNA duplexes (30ng/µl) were microinjected into the pronucleus/cytoplasm of the zygotes at the pronuclei stage. Injected zygotes were transferred into the uterus of pseudo pregnant F1 females. Genome-edited F1 *Gpr65*$^{-/-}$ were bred with C57BL/6 mice for two rounds to dilute the off-target effects. *Gpr65*$^{-/-}$ littermates were generated by breeding *Gpr65*$^{+/-}$ mice. *Gpr65*$^{gfp/gfp}$ mice were purchased from The Jackson Laboratory (stock number 008577). *Gpr65*$^{gfp/gfp}$ mice have 90% of exon 2 coding sequences replaced by promoterless IRES-EGFP sequences to disrupt GPR65 function.[4] *Gpr65*$^{gfp/+}$ mice were generated by crossing *Gpr65*$^{gfp/gfp}$ mice with WT C57BL/6J mice. *Rag1*$^{-/-}$ mice on a C57BL/6J background were obtained from the Clive and Vera Ramaciotti Laboratories, Walter and Eliza Hall Institute of Medical Research. All mice were maintained under specific pathogen free and controlled environmental conditions.

Murine diets used in this study include: i) Normal chow diet (Barastoc rat and mouse pellets; Ridley); ii) control AIN93G (Specialty Feeds); iii) a diet lacking resistant starches (low resistant starch or low fiber, LF, SF09-028; Specialty Feeds); iv) a diet rich in resistant starches (high resistant starch or high fiber, HF, SF11-025; Specialty Feeds). The major nutritional parameters are listed in Table S2. To eliminate gut microbiota, mice were treated with a combination of enrofloxacin (10 mg/kg body weight/ day) and amoxicillin with clavulanic acid (50 mg/kg body weight/ day) in drinking water for a week before the experimental procedures and maintained on this antibiotic cocktail until the end of the experiment.

**Tissue Collection**

Upon euthanasia by $CO_2$ asphyxiation, blood, and various tissues (heart, kidney, liver, gut, cecal content, spleen) were collected for further analyses. Samples were either snap frozen using liquid nitrogen and stored at -80 °C for DNA/RNA extractions later, stored in formalin before paraffin embedding for histopathology studies, or kept in cold phosphate buffer saline (PBS) to be used fresh for flow-cytometry studies.

**Large intestine intraluminal pH measurements**

The pH was measured in samples collected from caecum and colon using a pre-calibrated Thermo-scientific Orion Star A211 pH meter as previously described[5,6]. An Orion PerpHec ROSS Combination pH Micro Electrode (Thermo-scientific, diameter 3mm) was used. We ensured the sample covered the probe tip and a stable reading was acquired.

**Large intestine interstitial fluid pH measurements**

Intestinal interstitial fluid pH was measured using a protocol generously shared by Professor Dominik Muller (Max Delbrück Center for Molecular Medicine in the Helmholtz Association, Berlin, Germany), adapted from their publication.[7] An Orion PerpHec ROSS Combination pH Micro Electrode (Thermo-scientific, diameter 3mm) was used.

**Blood pressure measurement**

BP, heart rate, and activity were measured using radiotelemetry in untreated WT and *Gpr65*$^{-/-}$ female and male mice (n=4-6/group, total n=20 mice) at 8-10 weeks of age. Mice were anaesthetised for implantation of a radiotelemetry probe (TA11PA-C10,

Data Sciences International, USA) into the left carotid artery as described previously.[8] Mice recovered from surgery for at least 10 days. The probe was then magnetically switched on and data were recorded for 10 seconds every 10 minutes (Ponemah software, Data Sciences International, USA) for three days. Data were averaged to obtain the 24h means for each of the variables.

We also recorded systolic BP in untreated, sham and Ang II treated, and diet-treated WT and *Gpr65*$^{-/-}$ male mice noninvasively using tail-cuff in a CODA non-invasive blood pressure system (Kent Scientific Corporation). Mice were acclimatized for 3 days before recorded measurements, which included baseline and weekly measurements for 4 weeks. During these, 5 acclimatization cycles and 10 measurement cycles were recorded. At least 4 acceptable measurements were used to calculate the average systolic BP.

**Metabolic characterization**

Mice were placed in metabolic cages individually for 24-hours. Body weight, food intake, water intake, urine excretion, and feces excretion were measured. Body composition was analyzed by an EchoMRI 3-in-1 machine (Houston, TX).

**Saline challenge studies**

To examine if GPR65-deficiency diuretic and natriuretic responses, we performed a saline challenge test according to the protocol kindly shared by Prof. Alicia McDonough (Keck School of Medicine of University of Southern California).[9,10] Briefly, mice were intraperitoneally injected with 10% of their body weight of 37°C 0.9% saline solution and placed immediately in metabolic cages without any food or water. Urine volume and the amount of sodium excreted were measured hourly for 5 hours and are shown as both values per hour and accumulated over the 5-hour period. Sodium content was determined at the Department of Pathology, Monash Health.

**Cardiac ultrasound**

The day prior to the endpoint, mice were anesthetized using isoflurane and B-mode and M-mode echocardiography was performed to image the left ventricle using a Vevo 2100 Ultra High Frequency ultrasound system and a MS550D transducer by an experienced user at Monash Biomedical Imaging Centre (E.S.). Images were analyzed blind by L.X. and validated by an independent and experienced ultrasonographist (E.S.).

**Histological analysis**

Mouse heart, kidney, and gut tissues were fixed in 10% neutral buffered formalin for 24-48 hours, processed, paraffin embedded and sectioned at 4 μm. Masson's trichrome staining was performed on the sections to analyze the collagen present in heart, kidney, and intestinal samples. Periodic acid-Schiff alcian blue (PAS/AB) staining was performed on the gut sections to analyze goblet cells in the gut. Whole tissue sections were scanned using the Scanscope AT Turbo (Aperio) at 40× magnification. Total collagen in the heart and kidney tissue was quantified using a color thresholding macro using ImageJ software (FIJI).[11] Intestinal muscularis propria width, villi length, fibrosis and goblet cell frequency were quantified manually using FIJI ImageJ software, with each intestinal tissue section imaged three times in different fields of view. Magnifications for analyses are shown in the figure legends respectively.

**Isolation of mouse immune cells from spleen, kidney, and colon**

For mouse splenic cell suspensions, spleens were mechanically disrupted in cold PBS and passed through a 70μm strainer. Cells were then subjected to red blood cell lysis and washed with cold PBS.

For mouse renal cell suspensions, the kidneys were dissociated with scissors and digested in 1.5ml RPMI1640 with 0.156mg/ml Collagenase type XI, 0.030mg/ml Hyaluronidase, 1.8mg/ml Collagenase type I for 60 minutes at 37°C whilst shaking at 120rpm. Supernatant was decanted through a 70μm cell strainer then resuspended in 2ml of 40% isotonic Percoll solutions and underlaid with 2 ml of 80% isotonic Percoll solutions. Samples were centrifuged at 1400g for 25min at room temperature without using the brake. The layer of mononuclear cells at the interface were carefully collected and washed with PBS.

Mouse colon lamina propria cell suspension was prepared using a Lamina Propria Dissociation Kit, mouse (Miltenyi Biotec) following manufaturer's instruction. The cell pellets were then washed with PBS.

**$CD8^+$ T cells adoptive transfer**

To isolate $CD8^+$ T cells, single-cell suspensions of splenocytes were subject to Dynabeads™ Untouched™ Mouse CD8 Cells Kit (ThermoFisher Scientific) for the isolation of $CD8^+$ leukocytes according to the manufacturer's manuals. $5x10^6$ of either

WT or *Gpr65*$^{-/-}$ CD8$^{+}$ T cells were resuspended in 200uL PBS and intravenously injected in to *Rag1*$^{-/-}$ mice.

**Angiotensin-II induced hypertension model**

For hypertension experiments (except the adoptive transfer model), six-week-old male WT and *Gpr65*$^{-/-}$ mice received a subcutaneous minipump (Alzet model 2004) placed during surgery under anesthesia with isoflurane containing angiotensin II (Ang II, 0.5 mg/kg body weight/day; Auspep) for 4 weeks. Littermate mice were randomized into either group using an Excel randomization tool.

For the adoptive transfer model, 1 day after the adoptive transfer of the CD8$^{+}$ T cells, ten-week-old male *Rag1*$^{-/-}$ recipient mice underwent a subcutaneous minipump (Alzet model 1002) implantation containing Ang II at 1.44 mg/kg body weight/day (Auspep) for 2 weeks as described above. Littermate mice were randomized into either group using an Excel randomization tool.

**Flow cytometry**

Cells were resuspended with FACS buffer (PBS containing 2% FCS and 4 mM EDTA). Cell viability was evaluated using the fixable viability stain 620 (BD Horizon). After blocking of Fc receptors with mouse FcR blocking reagents (Miltenyi Biotech) in dark for 15 mins at room temperature, single cell suspension was surface stained in dark at 4°C for 30 min. The following antibodies were used for surface staining in T cell analysis panel: anti-mouse CD45 Alexa Fluor 700 (AF700; 30-F11; BD Pharmingen, San Jose, CA), anti-mouse TCRβ BUV496 (H57-597; BD Horizon), anti-mouse TCRγδ APC (GL3; BioLegend, San Diego, CA), anti-mouse CD4 BV510 (RM4-5; BD Horizon), anti-mouse CD8a FITC (53-6.7; BioLegend). The following antibodies were used for surface staining in myeloid cell analysis panel: anti-mouse CD45 Alexa Fluor 700 (AF700; 30-F11; BD Pharmingen, San Jose, CA), anti-mouse CD11b BV510 (M1/70, BioLegend), anti-mouse CD11c FITC (HL3; BD Pharmingen), anti-mouse Ly-6G BV421 (1A8; BD Horizon), anti-mouse F4/80 PE (T45-2342; BD Pharmingen), anti-mouse MHC-II PE-Cy7 (AF6-120.1; BioLegend). The following antibodies were used for surface staining in T cell phosphorylated CREB (pCREB) detection panel: anti-mouse CD4 BV510 (RM4-5; BD Horizon), anti-mouse CD8a AF700 (53-6.7; BD Pharmingen), anti-mouse TCRβ PE-Cy7 (H57-597; BD Pharmingen). The following antibodies were used for surface staining in T cell intracellular cytokine analysis panel: anti-mouse CD4 BV510 (RM4-5; BD Horizon), anti-mouse CD4 PE (GK1.5; BD Pharmingen), anti-mouse CD4 Pacific Blue (RM4-5; BD Pharmingen), anti-mouse CD8a AF700 (53-6.7; BD Pharmingen), anti-mouse TCRβ PE-Cy7 (H57-597; BD Pharmingen). The following antibodies were used for leukocyte detection in adoptive transfer experiment: anti-mouse CD45 AF700 (30-F11; BD Pharmingen), anti-mouse CD11b BV421 (M1/70, BioLegend), anti-mouse F4/80 PE (T45-2342; BD Pharmingen), anti-mouse TCRβ PE-Cy7 (H57-597; BD Pharmingen), anti-mouse CD8a FITC (53-6.7; BioLegend). Cells were then fixed and permeabilized in dark at 4°C using an Foxp3/Transcription Factor Staining Buffer Set (eBioscience). For $T_{reg}$ detection, fixed and permeabilized cells were then intracellular stained with anti-mouse Foxp3 PE (FJK-16s, eBiosciences) in dark at 4°C for 30 mins. For pCREB detection, cells were intracellular stained with rabbit anti-mouse pCREB (87G3, Cell Signaling) in dark at room temperature for 45 mins and subsequently stained with goat anti-rabbit IgG AF488 (Invitrogen) in dark at room temperature for 45 mins. For intracellular cytokine analyses, cells were intracellular stained with anti-mouse IFNγ BV421 (XMG1.2, BD Horizon), anti-mouse IFNγ eFluor450 (XMG1.2, eBioscience), anti-mouse TNFα FITC (MP6-XT22, BD Pharmingen), anti-mouse TNFα PE-Cy7 (MP6-XT22, Biolegend) in dark at 4°C for 30 mins. Cell counts were determined using flow cytometry counting beads (CountBright Absolute; Life Technologies) following manufacturer instructions. Sample data were acquired using a five-laser BD LSRFortessa X-20 flow cytometer and BD FACSDiva software (BD Biosciences) and analyzed using FlowJo software (Tree Star).

***In vitro* cell stimulation**

Mouse splenic or colon lamina propria cells were cultured in RPMI1640 supplemented with 10% FBS and 4 mM l-glutamine, which was altered to pH 6.5 or pH 7.5 using HCl and NaOH. Cells were stimulated with 100 ng/ml PMA (Sigma-Aldrich) and 1 μg/ml ionomycin (Sigma-Aldrich) for 2-h for pCREB detection, and for 4-h for intracellular cytokine analyses.

**cAMP assay**

$1\times10^6$ splenic cells were centrifuged and resuspended in 200 μL RPMI1640 supplemented with 10% FBS and 4 mM l-glutamine at pH 6.5 or pH 7.5. After incubating at 37°C for 30 mins, cells were washed with PBS at pH 6.5 or pH 7.5, respectively. Intracellular cAMP levels were measured by Invitrogen cAMP Competitive ELISA kit (ThermoFisher, #Cat: EMSCAMPL), following the manufacturer's instructions.

**DNA extraction from cecal contents and feces**

DNA extraction of cecal contents and feces was performed using the DNeasy PowerSoil DNA isolation kit (Qiagen, Germany) according to the manufacturer's protocol. DNA samples were quantified using Nanodrop (ThermoFisher Scientific).

**Mouse fecal bacterial load measurement**

Absolute bacterial load was measured using quantitative real-time PCR. Briefly, we used Fast SYBRgreen (Thermo Scientific) and 16S primers 1114F/1221R in a QuantStudio 7 qPCR Instrument (Thermo Scientific). Samples were loaded at 10ng/well, amplified in triplicates and compared to a standard containing $10^2$ to $10^{10}$ bacterial copies.

**16S ribosomal RNA sequencing**

V4 region of the 16S ribosomal RNA (rRNA) was targeted and amplified using primers previously published[12]. 240ng of the amplicon product per sample were pooled and purified. Sequencing was performed at the Australian Genomic Research Facility (AGRF, Melbourne, Australia) on MiSeq instrument (Illumina) to generate 300bp paired-end reads.

**16S rRNA bioinformatics analyses**

Sequencing followed our protocol recently published elsewhere. Sequenced data in the form of FASTQ files were obtained from the sequencing facility and passed initial quality control (QC) using FASTQC version 3. QIIME2 (version 2019-10) workflow was used for downstream analysis.[13] Raw reads were first trimmed (to ensure the quality of at least 50% of reads were >20 Phred score). Reads were then merged, dereplicated and chimeric reads were removed. The DADA2 QIIME2 plugin was used to 'denoise' the reads.[14] The SILVA database were used as a classifier for all taxonomy-related plots and tables.[15] The α-diversity rarefaction curves using the Shannon Index was performed to determine the ideal sequencing depth for further analyses. A minimum sampling depth of 10,000 annotated sequence variants (ASVs) was used unless stated otherwise.

β-diversity metrics as weighted Unifrac metrics were generated from the rarefied samples and shown as Principal Coordinate Analysis (PCoA) plots. Shannon, Faith's phylogenic diversity (Faith's PD) and Observed Operational Taxonomic Units (OTUs) were used to quantify α-diversity. The PERMANOVA (999 permutations for pseudo-F distribution) was used to test the separation between genotypes and diets. Significance was set as $P<0.05$, and results were adjusted for false discovery rate (FDR), where FDR $q<0.05$ was considered significant. Linear discriminant analysis (LDA) effect size (LEfSe)[16] was used to identify differentially abundant taxa between groups, with a specified effect size cut-off of 2.0 and Kruskal-Wallis test $P<0.05$. This data was validated using edgeR differential abundance analysis (false discovery rate adjusted $q<0.05$ on species) on MicrobiomeAnalyst.[17,18]

**Short-chain fatty acids quantification by liquid chromatography-mass spectrometry (LC-MS)**

Quantification of SCFAs including acetate, propionate, butyrate, isobutyrate, isovalerate, valerate and caproate was performed using LC-MS method, as previously described.[11,19] Briefly, 20μl of plasma and 20-40 mg of cecal content were analyzed in duplicates in a Q-Exactive Orbitrap mass spectrometer (Thermo Fisher Scientific) in conjunction with a Dionex Ultimate® 3000 RS high-performance liquid chromatography (HPLC) system (Thermo Fisher Scientific). We accepted a coefficient of variability <15%. Standard curves were constructed using the area ratio of the target analyte, and the internal standard in the range of each analyte was used. The levels of butyrate in some samples were below detection level of the standard curve; in this case they were considered half of the lowest measurable value, equivalent to 0.05 ng/ml.

#### Supplementary Tables

**Table S1. A phenome-wide association study summary for *GPR65* using the Atlas of GWAS Summary Statistics**.

[see full table attached in Excel]

Legend: N: sample size.

**Table S2. Calculated nutritional parameters of murine diets used in this study.**

| | Barastoc Mice Breeder Cube | AIN93G | SF09-028 Low Fiber | SF11-025 High Fiber* |
|---|---|---|---|---|
| **Crude Fiber** | 3.2% | 4.7% | 0.0% | 9.7% |
| **Crude Protein** | 20% | 19.7% | 19.4% | 19.4% |
| **Crude Fat** | 8.5% | 7.0% | 7.0% | 7% |
| **Acid Detergent Fiber** | 4.4% | 4.7% | 0.0% | 9.7% |
| **Digestible Energy** | 13.2 MJ/kg | 16.2 MJ/kg | 16.9 MJ/kg | 15.6 MJ/kg |
| **Calcium** | 1.1% | 0.69% | 0.47% | 0.47% |
| **Phosphorus** | 0.96% | 0.35% | 0.32% | 0.32% |
| **Sodium** | 0.35% | 0.16% | 0.12% | 0.12% |
| **Potassium** | 0.89% | 0.40% | 0.40% | 0.40% |
| **Chloride** | 0.57% | 0.16% | 0.16% | 0.16% |
| **Magnesium** | 0.25% | 0.07% | 0.09% | 0.10% |
| **Lysine** | 1.22% | 1.64% | 1.50% | 1.50% |
| **Methionine** | 0.38% | 0.92% | 0.80% | 0.80% |
| **Iron** | 180 mg/kg | 49 mg/kg | 73 mg/kg | 75 mg/kg |
| **Zinc** | 103 mg/kg | 46 mg/kg | 53 mg/kg | 53 mg/kg |
| **Manganese** | 156 mg/kg | 16 mg/kg | 18 mg/kg | 18 mg/kg |
| **Copper** | 20 mg/kg | 7.0 mg/kg | 7.1 mg/kg | 7.3 mg/kg |
| **Molybdenum** | 1.19 mg/kg | 0.15 mg/kg | 0.15 mg/kg | 0.15 mg/kg |
| **Iodine** | 1.67 mg/kg | 0.2 mg/kg | 0.2 mg/kg | 0.2 mg/kg |
| **Linoleic Acid** | 3.11% | 2.49% | 2.49% | 2.48% |
| **Selenium** | 0.33 mg/kg | 0.2 mg/kg | 0.3 mg/kg | 0.3 mg/kg |
| **Cobalt** | 0.67 mg/kg | No data | No data | No data |
| **Vitamin A** | 15 IU/g | 4 IU/g | 4 IU/g | 4 IU/g |
| **Vitamin D** | 2 IU/g | 1 IU/g | 1 IU/g | 1 IU/g |
| **Vitamin E** | 270 mg/kg | 78 mg/kg | 78 mg/kg | 78 mg/kg |
| **Vitamin K** | 59 mg/kg | 1 mg/kg | 1 mg/kg | 1 mg/kg |
| **Vitamin B1** | 69 mg/kg | 6 mg/kg | 6.1 mg/kg | 6.1 mg/kg |

| | | | | |
|---|---|---|---|---|
| **Vitamin B2** | 52 mg/kg | 6.4 mg/kg | 6.3 mg/kg | 6.3 mg/kg |
| **Vitamin B6** | 34 mg/kg | 7 mg/kg | 7 mg/kg | 7 mg/kg |
| **Vitamin B12** | 80 µg/kg | 100 µg/kg | 103 µg/kg | 103 µg/kg |
| **Niacin** | 437 mg/kg | 30.1 mg/kg | 30 mg/kg | 30 mg/kg |
| **Pantothenic Acid** | 235 mg/kg | 16 mg/kg | 16.5 mg/kg | 16.5 mg/kg |
| **Biotin** | 1.68 mg/kg | 0.2 mg/kg | 0.2 mg/kg | 0.2 mg/kg |
| **Folic Acid** | 11.82 mg/kg | 2 mg/kg | 2 mg/kg | 2 mg/kg |

*All carbohydrate has been replaced with Gel Crisp starch, which is a modified type 2 resistant, acetylated high amylose resistant starch made from maize starch.

#### Supplementary Figures

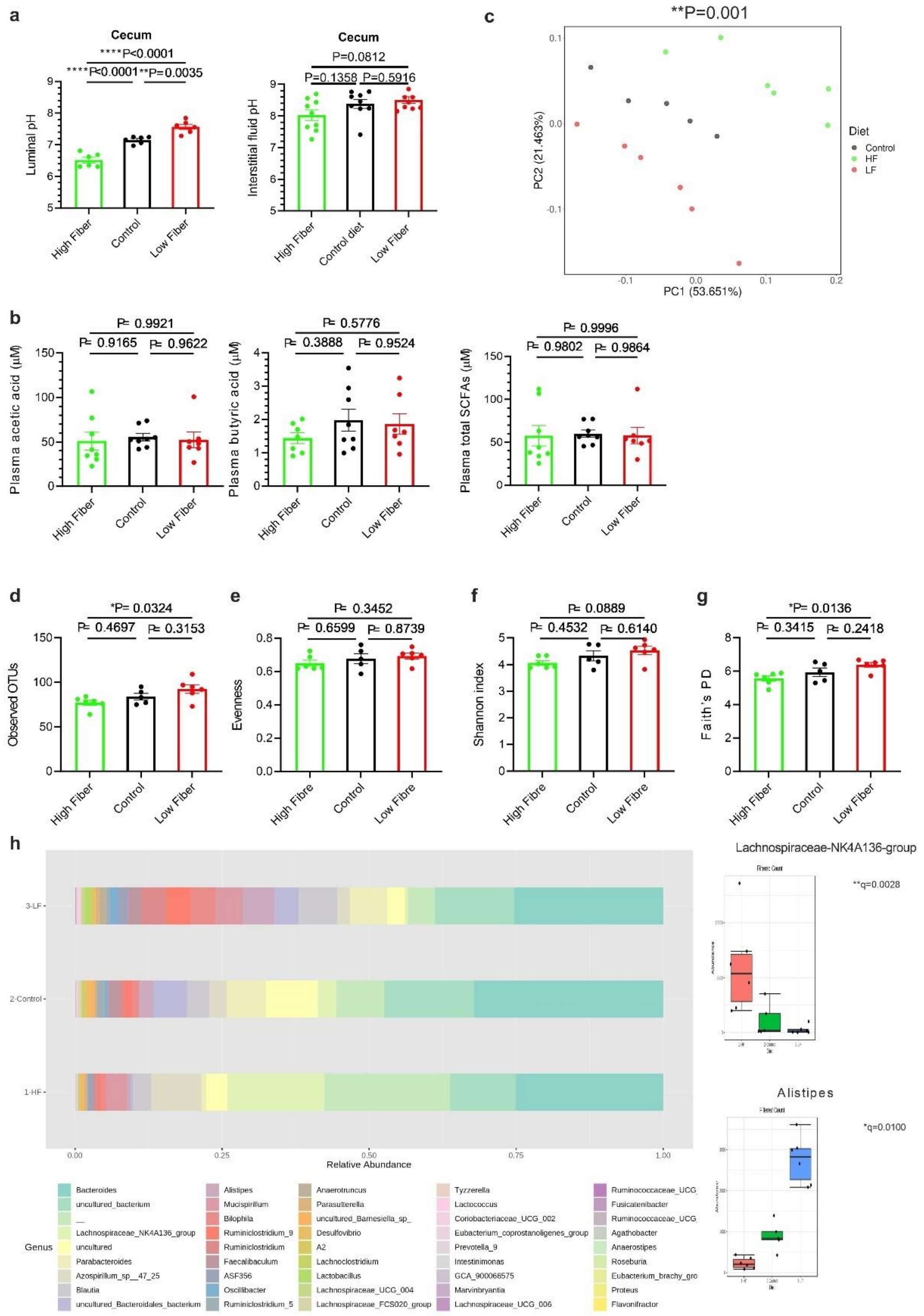


**Figure S1. Dietary fiber significantly alters the gut microbiome which is responsible for the pH regulations.**

**A**, Cecum content pH (left), but not cecum interstitial fluid pH (right), changes according to diet. **B,** Levels of acetic and butyric acids and total SCFAs in plasma from the peripheral circulation (cardiac puncture). **C-I**, Gut microbiome (V4 region of the 16S rRNA) changes according to diet, showing markers of **C**, β-diversity shown as a weighted Unifrac principal coordinate analysis and **D-G,** α-diversity indicated by observed operational taxonomic units (OTUs), evenness, Shannon's index, and Faith's PD. **H,** Major changes in the microbiome according to diet. **I**, Differential abundance of *Lachnospiraceae*-NK4A136-group and *Alistipes* spp. Each data point represents an individual mouse, and n=6 mice in each group; All data represented as means ± SEM; n=5-8 mice in each group; **A**, **B, D-F**, One-way ANOVA with adjustment by false discovery rate; **C**, PERMANOVA; **G-H**, EdgeR algorithm. Legend: HF, high fiber; LF, low fiber.

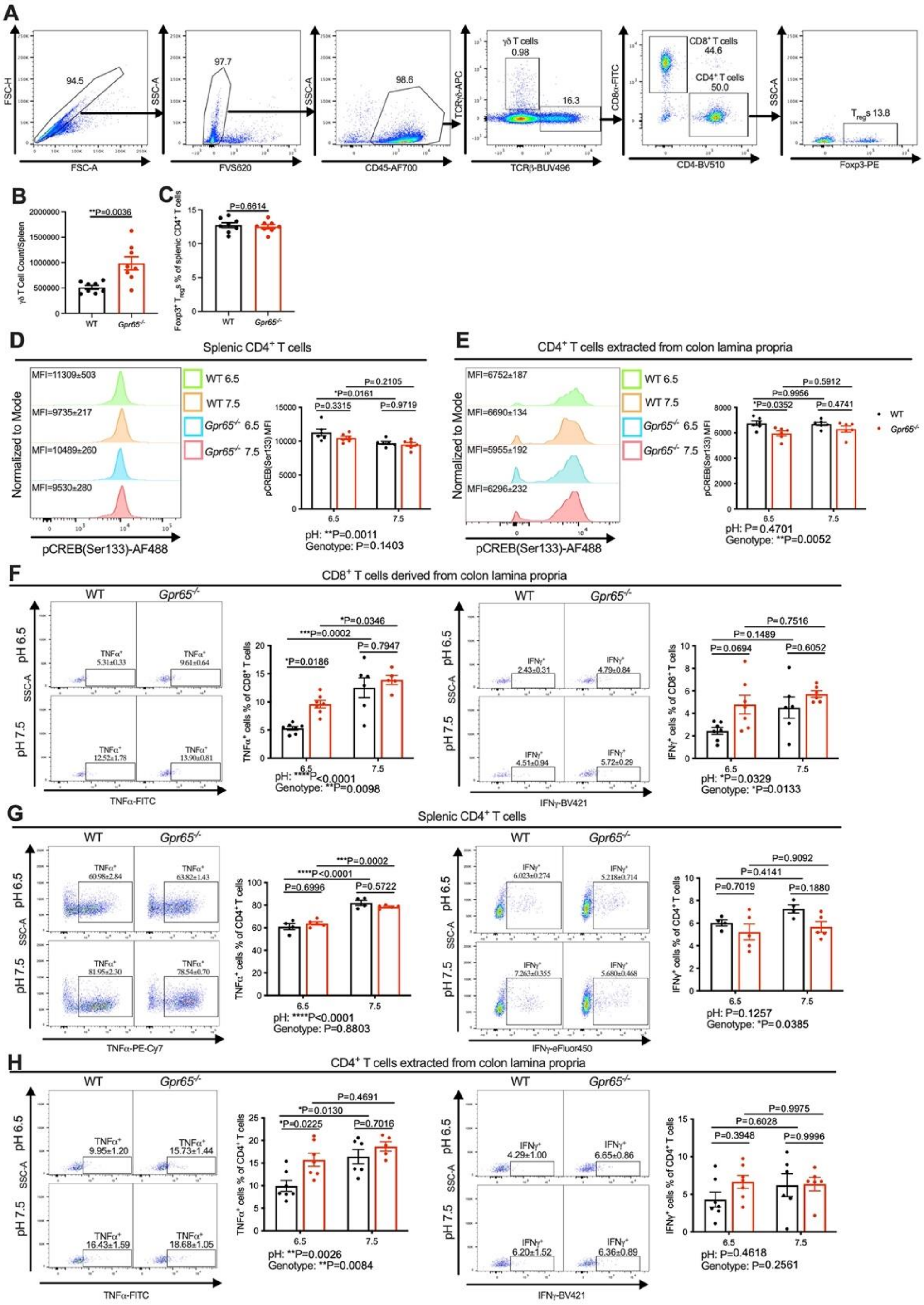


**Figure S2. The impact of pH and GPR65 signaling on cAMP-pCREB-cytokine production axis.**

Splenocytes were isolated from mice at the age of 12 weeks. **A**, Gating strategy for T cell analysis; **B**, γδ T cell count in the spleen. **C**, Percentage of Foxp3$^{+}$ T$_{reg}$s from splenic CD4$^{+}$ T cells. 1×10$^{6}$ splenocytes and colon lamina propria cells were incubated with 100ng/ml PMA and 1μg/ml ionomycin for 2 hours. **D**, pCREB (Ser133) levels of splenic CD4$^{+}$ T cells. **E**, pCREB (Ser133) levels of CD4$^{+}$ T cells extracted from colon lamina propria. 1×$10^6$ splenocytes and colon lamina propria cells were incubated with 100ng/ml PMA and 1μg/ml ionomycin for 4 hours. **F**, Percentage of TNFα and IFNγ producing $CD8^+$ T cells at pH 6.5 and 7.5 among WT and *Gpr65*$^{-/-}$ $CD8^+$ T cells derived from colon lamina propria. **G**, Percentage of TNFα and IFNγ producing $CD4^+$ T cells at pH 6.5 and 7.5 among WT and *Gpr65*$^{-/-}$ splenic $CD4^+$ T cells. **H**, Percentage of TNFα and IFNγ producing $CD4^+$ T cells at pH 6.5 and 7.5 among WT and *Gpr65*$^{-/-}$ $CD4^+$ T cells derived from colon lamina propria. Each data point represents an individual mouse, and n=4-9 mice in each group; All data represented as means ± SEM. **B** and **C**, Student's t test. **D-H**, Two-way ANOVA. *P<0.05, **P<0.01, ***P<0.001, ****P<0.0001.

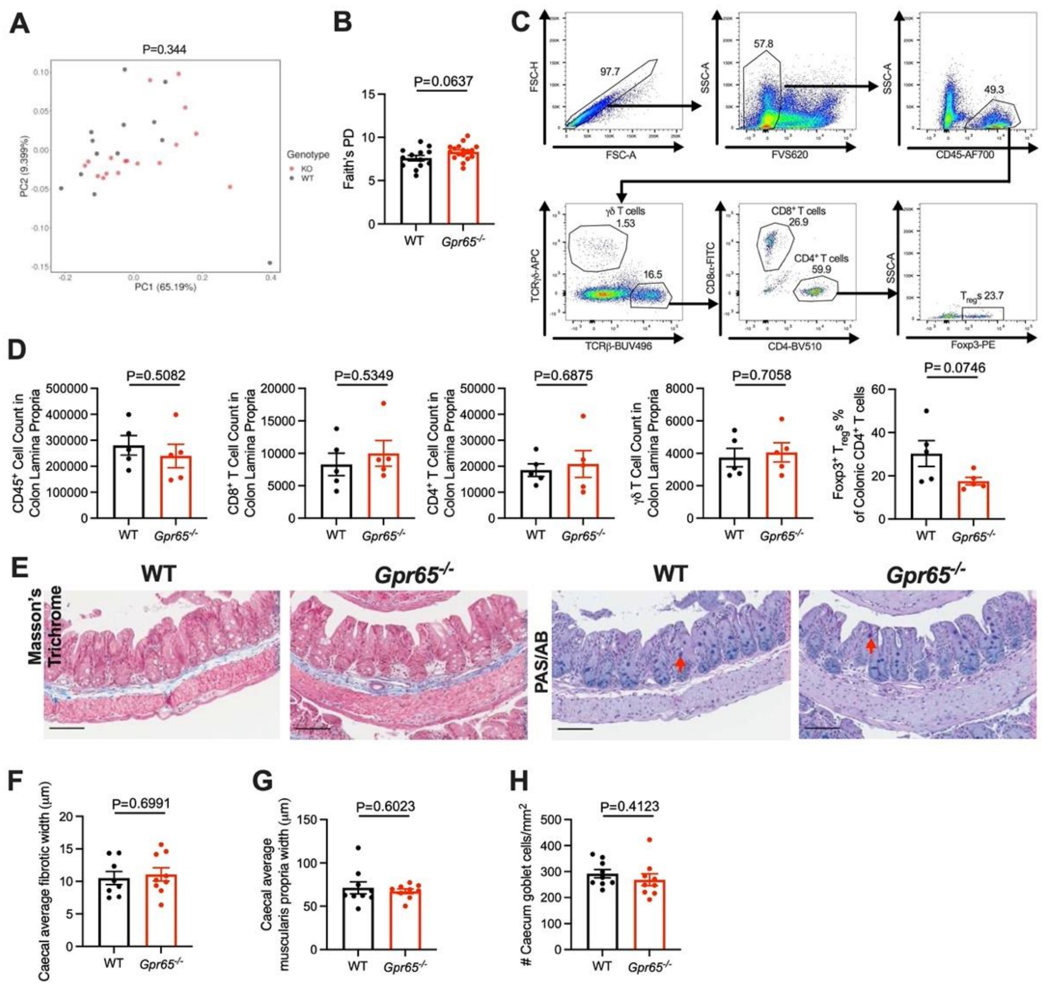


**Figure S3. GPR65 deficiency does not result in spontaneous differences in gut homeostasis.**

DNA was extracted from mouse cecal contents collected. V4 region of the 16s rRNA was sequenced. **A**, β-diversity shown as a weighted Unifrac principal coordinate analysis. **B**, α-diversity indicated by Faith's phylogenic diversity index (Faith's PD). Single cell suspensions were isolated from mouse colon lamina propria at the age of 12 weeks. **C**, Gating strategy; **D**, $CD45^+$ immune cell count, $CD8^+$ T cell, $CD4^+$ T cell, γδ T cell count in colon lamina propria, and percentage of $Foxp3^+$ $T_{reg}$s from colonic $CD4^+$ T cells. Cecum tissues were collected and cecal sections were stained with Masson's trichrome or Periodic acid-Schiff alcian blue (PAS/AB). **E**, Representative caecum sections, magnification = ×100, scale bar = 100μm, Red arrows point to examples of goblet cells; **F**, Average fibrotic width; **G**, Average muscularis propria width; **H**, Goblet cell frequency. Samples were collected at the age of 12 weeks **A**, n=12-17 mice in each group; PERMANOVA. **B**, **D**, **F-H**, each data point represents an individual mouse, and n=7-17 mice in each group; All data represented as means ± SEM; Student's t test. Legend: KO, GPR65 knockout mouse; WT, wild-type mouse.

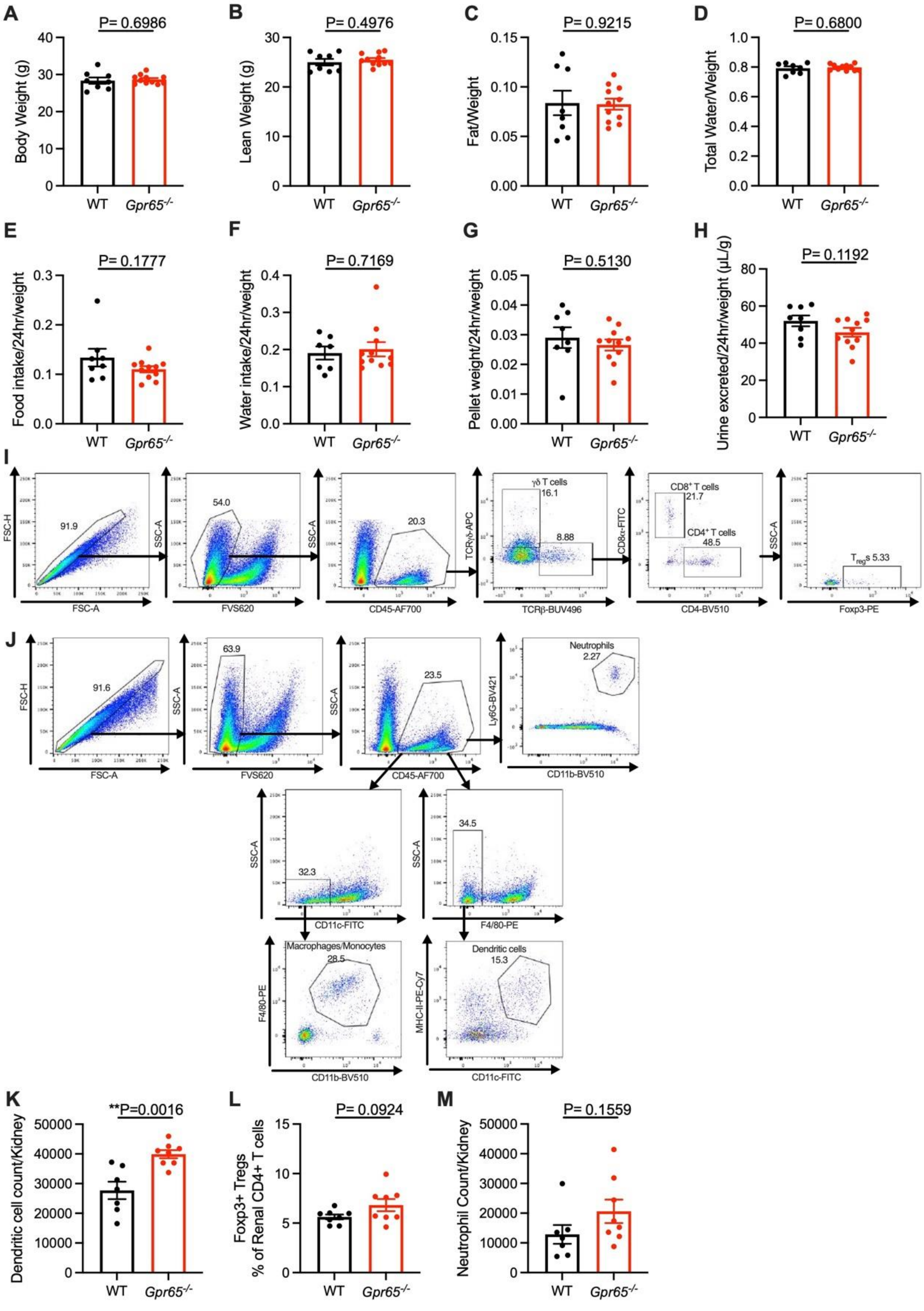


**Figure S4. The impact of GPR65 deficiency on metabolic phenotypes and renal immune cell infiltration in male mice.**

EchoMRI and metabolic cage results of male mice performed at the age of 10 weeks. **A**, Body weight; **B**, Lean weight; **C**, Fat weight/body weight ratio; **D**, Total water weight/body weight ratio. 24h metabolic cage of male mice was performed at the age of 11 weeks. **E**, 24h food intake adjusted by body weight; **F**, 24h water intake adjusted by body weight; **G**, 24h faeces excretion adjusted by body weight; **H**, 24h urine excretion adjusted by body weight. Single cell suspensions were prepared from kidneys of male mice at the age of 12 weeks. **I**, Gating strategy for T cell analysis; **J**, Gating strategy for myeloid cell analysis; **K**, Dendritic cell count; **L**, Percentage of $Foxp3^+$ $T_{reg}$s from renal $CD4^+$ T cells; **M**, Neutrophil count. Each data point represents an individual ear or mouse, and n=6-11 mice in each group; All data represented as means ± SEM; Student's t test. **P<0.01. Legend: WT, wild-type mice; FVS620, fixable viability stain 620.

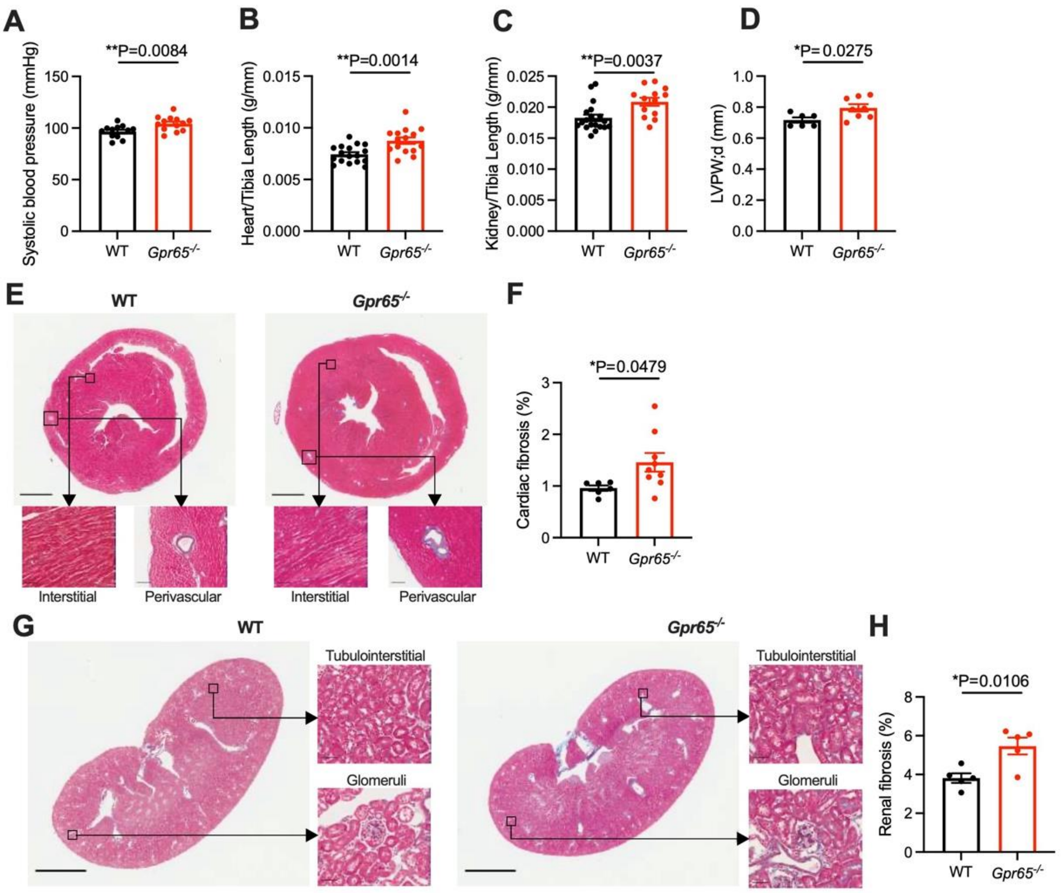


**Figure S6. GPR65 deficiency spontaneously increases blood pressure and tissue hypertrophy in female mice.**

Blood pressure (BP) of female mice was measured non-invasively by tail cuff at the age of 10 weeks. **A**, Systolic BP. Tissues of female mice were collected and weighed at the age of 12 weeks. **B**, Heart weight to tibia length index. **C**, Kidney weight to tibia length index. Cardiac ultrasound screening was performed at the age of 11 weeks. **D**, Left ventricle posterior wall in the end diastole. Heart sections were stained with Masson's trichrome. **E**, Representative heart sections; For upper panels, magnification = ×10, scale bar = 1mm; For lower left panels (interstitial), magnification = ×200, scale bar = 50μm; For lower right panels (perivascular), magnification = ×100, scale bar = 100μm. **F**, Percentage of fibrotic area in the cardiac tissues. Kidney sections were stained with Masson's trichrome. **G**, Representative kidney sections; For left panels, magnification= ×5, scale bar = 2mm; For right upper panels (tubulointerstitial), magnification = ×200, scale bar = 50μm; For right lower panels (glomeruli), magnification = ×200, scale bar = 50μm. **H**, Percentage of fibrotic area in the renal fibrosis. Each data point represents an individual mouse, and n=6-19 mice in each group; All data represented as mean ± SEM; Student's t test. *P<0.05, **P<0.01. Legend: WT, wild-type mice.

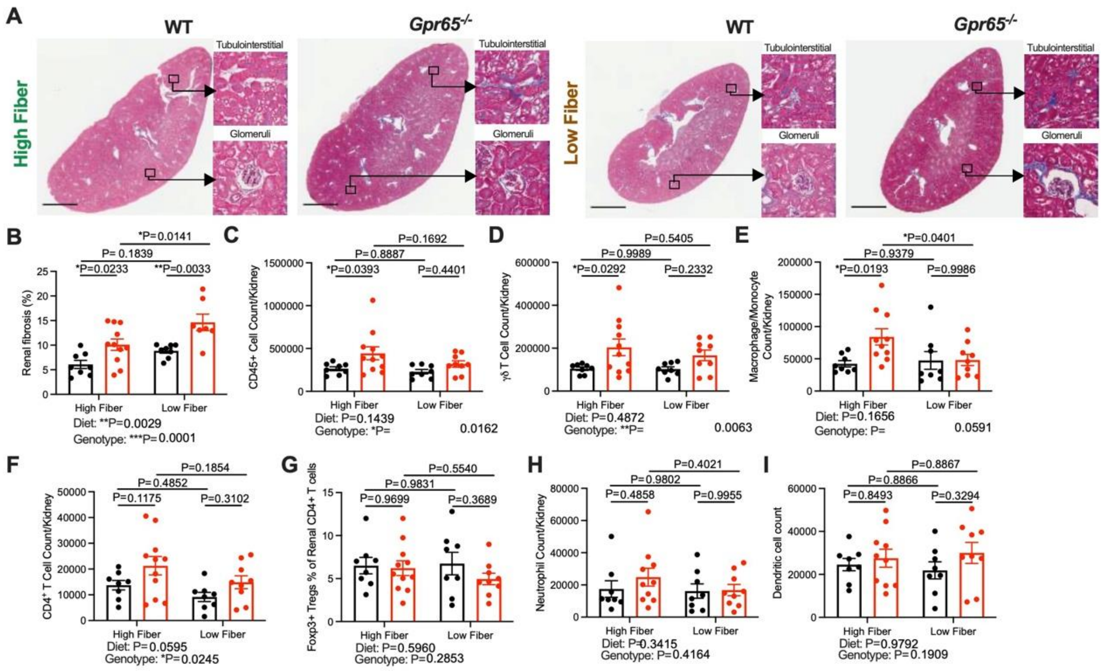


**Figure S6. The impact of dietary fiber and GPR65 on BP and renal function and immune cell infiltration under hypertensive stimuli.**

6-week-old male WT and *Gpr65*$^{-/-}$ were infused with 0.5mg/kg body weight/day angiotensin-II for 28 days and fed with either high fiber (HF) or low fiber (LF) diet. **A**, Representative kidney sections; For left panels, magnification= ×5, scale bar = 2mm; For right panels, magnification = ×200, scale bar = 50µm. **B**, Percentage of fibrotic area in the renal tissues. Mice were intraperitoneally injected with 10% of their body weight of 37 °C 0.9% saline solution at the 3$^{rd}$ week post minipump implantation. Single cell suspensions were prepared from kidneys of the mice at the endpoint of the study. **C**, CD45$^{+}$ immune cell count; **D**, γδ T cell count; **E**, Macrophage/Monocyte count; **F**, CD4$^{+}$ T cell count; **G**, Percentage of Foxp3$^{+}$ $T_{reg}$s from CD4$^{+}$ T cells; **H**, Neutrophil count; **I**, Dendritic cell count. Each data point represents an individual mouse, and n=7-13 mice in each group; All data represented as mean ± SEM. Two-way ANOVA. *P<0.05, **P<0.01, ***P<0.001, ****P<0.0001. Legend: WT, wild-type mice.

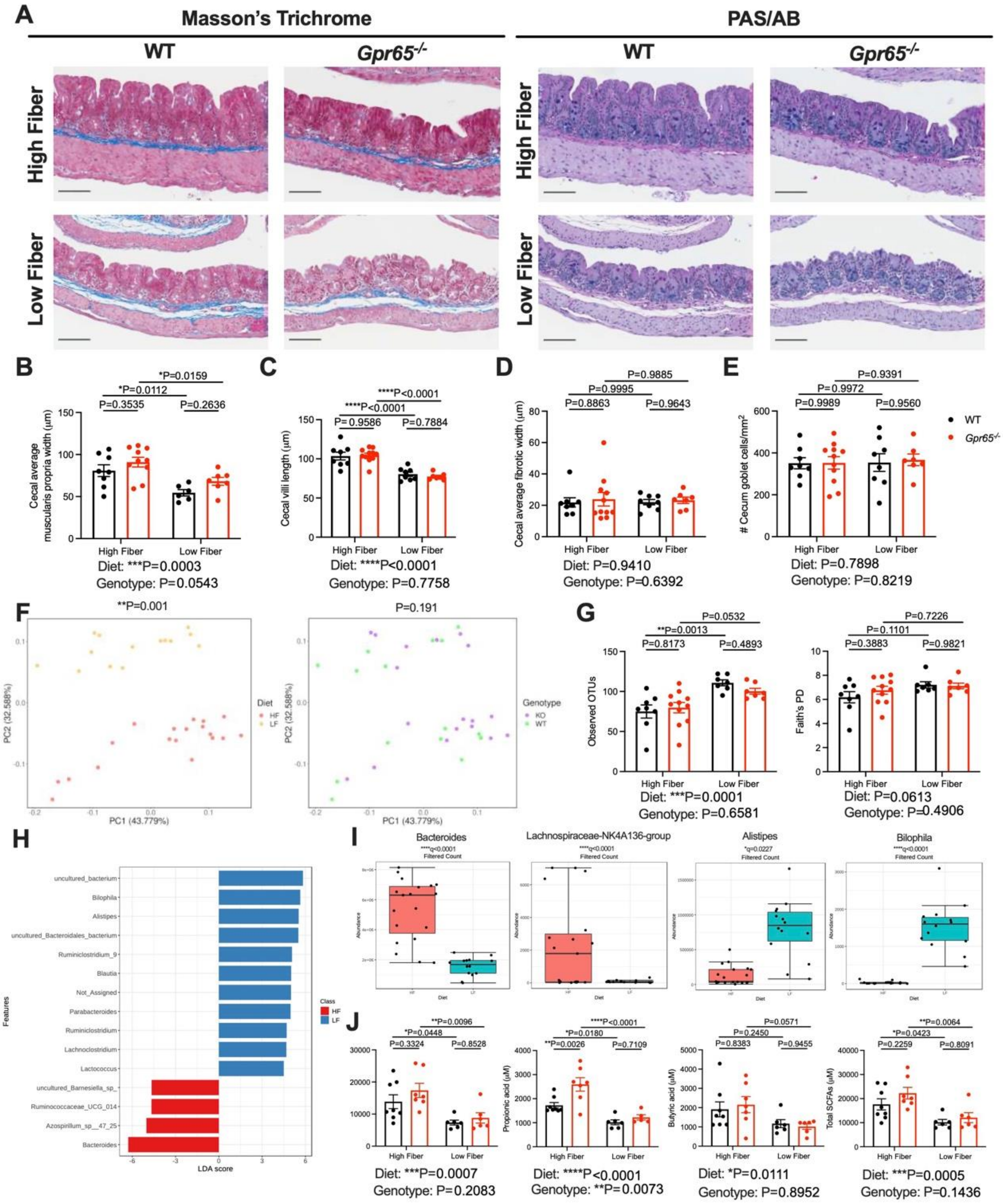


**Figure S7. The impact of dietary fiber and GPR65 on gut pathophysiology and microbiome composition under hypertensive stimuli.**

Cecum tissues were collected at the endpoint of the study and cecal sections were stained with Masson's trichrome or Periodic acid-Schiff alcian blue (PAS/AB). **A**, Representative caecum sections, magnification = ×100, scale bar = 100µm; **B**, Cecal average muscularis propria width; **C**, Cecal villi length; **D**, Cecal average fibrotic width; **E**, Goblet cell frequency in cecum; DNA was extracted from mouse cecal contents collected at the endpoint of the study. V4 region of the 16s rRNA was amplified and sequenced. **F**, β-diversity shown as a weighted Unifrac principal coordinate analysis. **G**, α-diversity indicated by observed operational taxonomic units (OTUs) and Faith's PD. **H**, Features at the genus level determining the differences of gut microbiota between mice fed with high fiber (HF) and low fiber (LF) diet. **I**, Abundance of *Bacteroides* spp., *Lachnospiraceae*-NK4A136-group, *Bilophila* spp., *Alistipes* spp.. **J**, Levels of acetic, propionic, butyric acids and total short-chain fatty acids (SCFAs) in cecal contents. Each data point represents an individual mouse, and n=5-11 mice in each group; All data represented as mean ± SEM. **B-E**, **G**, **J**, Two-way ANOVA; **F**, PERMANOVA; **I**, EdgeR algorithm. *P<0.05, ***P<0.001, ****P<0.0001. Legend: WT, wild-type mice.

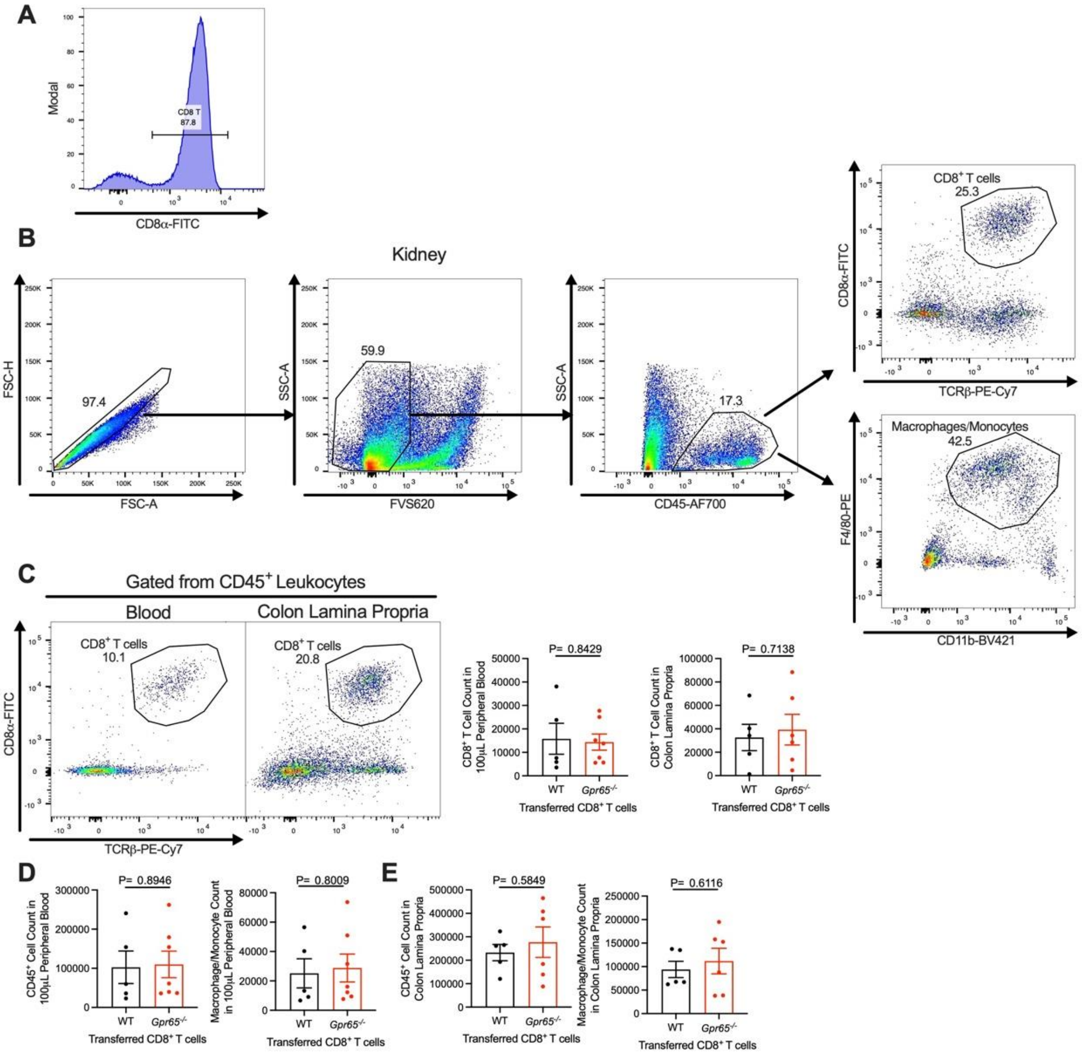


**Figure S8. Adoptive transfer of CD8$^+$ T cells.**

CD8$^+$ T cells were purified from the spleens of WT and *Gpr65*$^{-/-}$ mice. **A**, The purity of the CD8$^+$ T cells for adoptive transfer. 10-week-old male *Rag1*$^{-/-}$ were adoptive transferred with either WT and *Gpr65*$^{-/-}$ CD8$^+$ T cells, and then infused with 1.44 mg/kg body weight/day angiotensin-II for 14 days. They were fed with HF diet. Single cell suspensions were prepared from kidneys, blood, and colon lamina propria of the mice at the endpoint of the study. **B**, Gating strategy on the kidneys; **C**, CD8$^+$ T cell presence and count in 100µL blood and colon lamina propria; **D**, CD45$^+$ immune cell count and Macrophage/Monocyte count in 100µL blood; **E**, CD45$^+$ immune cell count and Macrophage/Monocyte count in colon lamina propria. Each data point represents an individual mouse, and n=5-7 mice in each group; All data represented as means ± SEM. Two-way ANOVA. *P<0.05, ****P<0.0001.
